# The interplay between metabolic stochasticity and regulation in single *E. coli* cells

**DOI:** 10.1101/2022.08.29.505271

**Authors:** Martijn Wehrens, Laurens H.J. Krah, Benjamin D. Towbin, Rutger Hermsen, Sander J. Tans

**Affiliations:** Hubrecht Institute, Royal Netherlands Academy of Arts and Sciences (KNAW) and University Medical Center, Utrecht, the Netherlands; Utrecht University, Theoretical Biology, Utrecht, The Netherlands; Institute of cell biology, University of Bern, Bern, Switzerland; AMOLF, Biophysics, Amsterdam, The Netherlands

## Abstract

Metabolism is inherently stochastic at the cellular level. Whether cells actively regulate processes in response to these random internal variations is a fundamental problem that remains unaddressed, yet critical to understanding biological homeostasis. Here, we show that in E. coli cells, expression of the main catabolic enzymes is continuously adjusted in response to metabolic fluctuations under constant external conditions. This noise feedback is performed by the cAMP-CRP system, which controls transcription of the catabolic enzymes by modulating concentrations of the second messenger cAMP upon changes in metabolite abundance. Using time-lapse microscopy, genetic constructs that selectively disable cAMP-CRP noise feedback, and mathematical modelling, we show how fluctuations circulate through this hybrid metabolic-genetic network at sub cell-cycle timescales. This circulation of stochastic fluctuations is explained by four distinct noise propagation modes, one of which describes the continuous cAMP-CRP regulation. The model successfully predicts how noise circulation is impacted by C-sector under and over-expression. The results raise the question whether the cAMP-CRP system, as well as other metabolic regulation mechanisms, have evolved to manage internal metabolic fluctuations in addition to external growth conditions. We conjecture that second messengers may broadly function to control metabolic stochasticity and achieve cellular homeostasis.

## Introduction

Bacteria display a striking ability to adapt to diverse environments. When exposed to different carbon sources, bacterial cells make vast changes to their proteome composition, allowing them to optimize their allocation of metabolic resources^1–4^. Many regulation mechanisms have been identified that adjust enzyme expression to the growth medium^5–8^. In addition to these external changes, however, bacteria are also confronted with major internal variations^9,10^. Gene expression has long been known to be stochastic^9–12^. The metabolic activity of cells was more recently found to fluctuate randomly in time under constant external conditions, thus severely limiting growth^13–17^. These observations raise the question whether cells also adjust their proteome to internal metabolic fluctuations, which differ fundamentally from external changes in growth media. Addressing this issue is key to understanding the elementary principles of cellular homeostasis and the functional relevance of known regulatory interactions.

Here, we address these issues using cAMP-CRP signaling in *Escherichia coli* as a model system. cAMP-CRP signaling is a major regulation mechanism of metabolic activity (Fig. 1A). Regulating over 180 genes, CRP is a general expression activator of a group of catabolic enzymes that together are referred to as the C-sector^4,18–20^. CRP is activated by the second messenger cyclic AMP (cAMP), whose synthesis is inhibited by metabolites that are produced by the C-sector enzymes. This negative cAMP-CRP feedback loop has been shown to produce a near-linear relation between the C-sector proteome mass fraction (*φ*_C_) and the growth rate (*λ*) under variation of the carbon source available in the medium^3,18^, and to optimally balance the costs and benefits of C-sector expression such that the overall growth rate is maximized in a range of nutrient conditions^5^. However, the interplay between internal stochastic variations in metabolic activity and the cAMP-CRP system or any other metabolic regulatory feed-back mechanism has not been addressed experimentally. Indeed, while the stochasticity of metabolic activity has been evidenced by correlations between cellular growth and enzyme expression noise^13^, the nature of these metabolic fluctuations is unclear, and different from the metabolic changes induced by external conditions. For instance, it is unknown which pathways are affected, which metabolites fluctuate in abundance, whether more global changes in proteome expression are involved, and indeed how the resulting combination of internal variations impact the cAMP-CRP system. This issue remains unresolved for any metabolic regulation mechanism, and hence addressing it for cAMP-CRP is a key first step in understanding the regulatory basis of metabolic homeostasis in the context of intrinsic molecular stochasticity.

**Figure 1.**
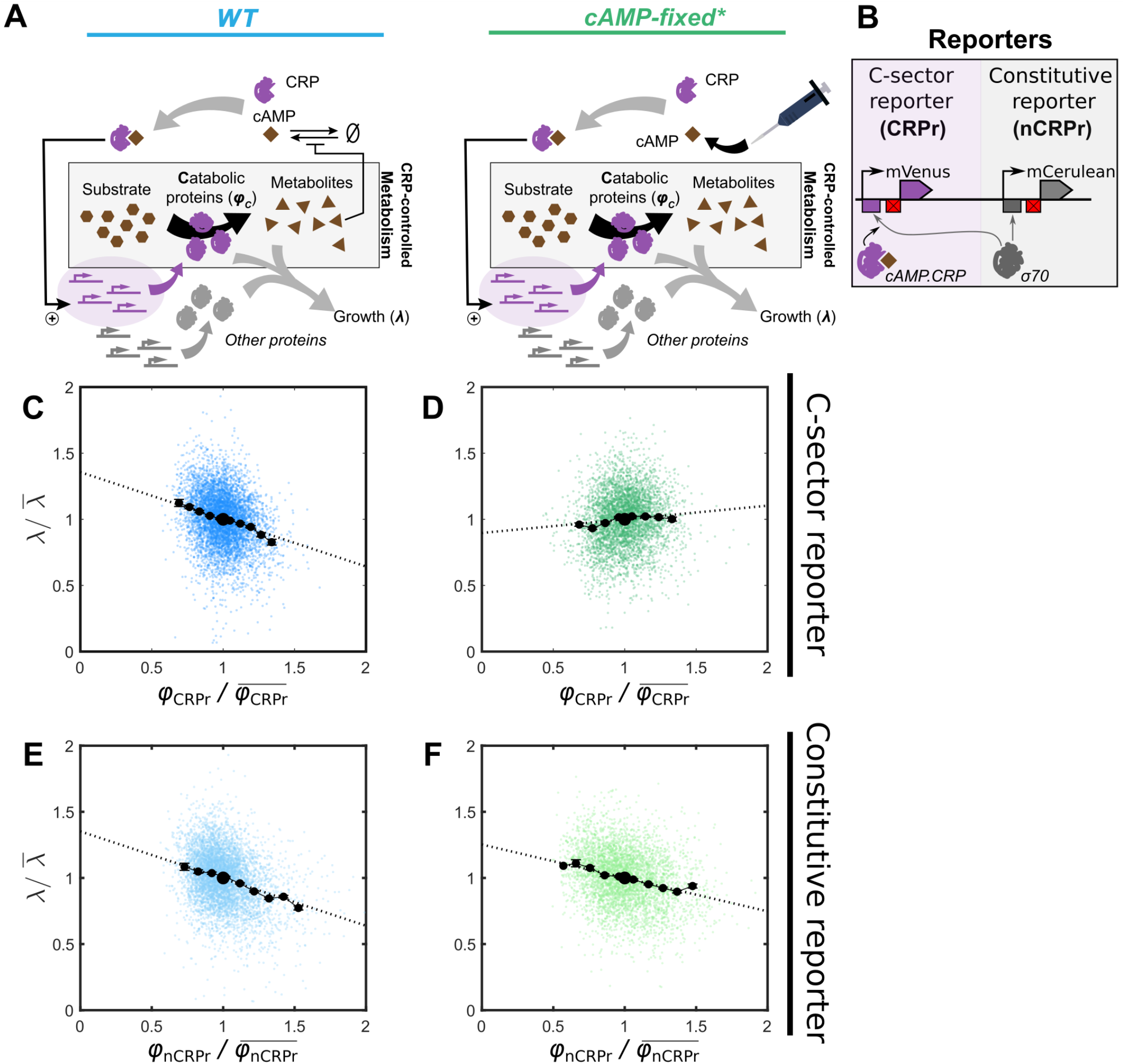
Removing cAMP feedback alters dynamics of a regulated reporter only. (A) Cartoon of a bacterial cell and the difference between wild type and mutant. Shown are the processes of metabolism, protein expression, cAMP-CRP regulation, and growth (λ). Here φ_C_ represents the expression level of the C-sector; the total concentration of all catabolic proteins that are regulated by cAMP-CRP and that import nutrients and convert them into internal metabolites (including cAMP itself). In the cAMP-fixed strain, cAMP is neither synthesized nor degraded, and instead supplied externally to experimentally tune φ_C_. (B) The two reporters and their promoters that were used in this study: a C-sector reporter whose transcription was regulated by cAMP-CRP (CRPr), and a constitutive reporter, nCRPr. Crossed red block is the scrambled lacI site. (C-F) Scatter plot of instantaneous growth rate (λ) against single-cell relative expression of the reporters (φ_CRPr_ or φ_nCRPr_). Dashed lines are linear regressions, black dots indicate binned averages. Plots are from single, representative micro-colonies (n = 1671 cells for WT, n= 1580 cells for cAMP-fixed*), other colonies showed the same trends, Fig S5).

To study stochasticity in the cAMP-CRP system we quantified the correlations between C-sector expression and growth rate in individual cells over time using time lapse microscopy, which allows identification of time delays in the propagation of fluctuating signals. To dissect the role of the cAMP-CRP feedback, we aimed to specifically disrupt its transmission of noise. The challenge here was to maintain the C-sector stimulation by cAMP, which is essential to growth, while inhibiting the feed-back of cAMP noise. To achieve this, we used an *E. coli* c*yaA cpdA* null mutant that is unable to synthesize or hydrolyze cAMP^5^. The noise feedback was thus broken while externally supplied cAMP allowed one to maintain C-sector expression at appropriate levels (Fig. 1A). In *WT* cells, C-sector expression showed fluctuations that were negatively correlated and time-delayed with respect to growth fluctuations, in line with cAMP-CRP regulating the C-sector in response to metabolic stochasticity. Consistently, disrupting the cAMP-CRP feedback abolished these negative correlations. A mathematical model we developed could explain all the observed correlations in a single fitting procedure, reproducing the observed effect of the disrupted feedback by merely changing the feedback coupling parameter to zero. This mechanistic understanding of the system was further evidenced by the ability of the model to predict the changes in stochastic dynamics for cAMP levels below and above wild type levels. Together, the findings show that C-sector expression in *E. coli* is continuously regulated in response to internal stochastic metabolic fluctuations in fixed environments. They also suggest that feed-backs in metabolic networks, which are ubiquitous in cells, act more generally to control and exploit internal metabolic noise.

## Results

### Interrupting the feedback of noise in the cAMP-CRP system

Elucidating noise propagation in the cAMP-CRP system requires insight into the stochasticity of the C-sector expression that is regulated by CRP. The mean C-sector population expression was previously studied by quantifying the expression of a representative enzyme, LacZ ^18^. Here we follow this general approach and measure the expression of mVenus driven by the *lac* promotor (Fig. 1B). As we aim to study fluctuations propagated by CRP rather than by the *lac* repressor LacI, the LacI binding site in the lac promotor is scrambled such that LacI no longer binds, while the promotor remains sensitive to cAMP-CRP^5,21^. This genome-inserted construct is called the CRP-regulated reporter, or CRPr. To study noise unrelated to CRP or LacI, a second reporter was created by further modifying the CRPr promoter. Specifically, the region where CRP and *σ*^70^ bind was replaced with a *σ*^70^ consensus site, such that transcriptional initiation occurs constitutively without requiring CRP to recruit *σ*^70^. This reporter construct, which was fused to mCerulean, is referred to as the non-CRP regulated reporter, or nCRPr (Fig. 1B, S11). Both the CRPr and nCRPr reporters were chromosomally inserted into *E. coli*, a construct we refer to as *WT*, and in a c*yaA cpdA* null mutant, which we refer to as *cAMP-fixed*.

Bulk measurements in lactose minimal media showed that the growth rate of *cAMP-fixed* peaked at about 800 μM externally supplied cAMP, while decreasing to almost negligible growth at lower and higher cAMP concentrations^5,22^ (Fig. S3). This strong dependence on cAMP is consistent with the many genes controlled by CRP and their essential nature. At low cAMP, under-stimulation of C-sector expression leads to decreased metabolic flux and concomitant growth. Conversely, at high cAMP, over-stimulation of C-sector expression leads to excess expression of many genes, which is metabolically costly and in turn also reduces the growth rate. We find that the growth rate of *WT* cells in the same lactose minimal medium (without externally supplied cAMP) is similar as the optimal growth rate of *cAMP-fixed* cells obtained at 800 μM cAMP (Figs. S3, S5B). We therefore refer to *cAMP-fixed* cells growing at 800 μM cAMP as *cAMP-fixed*\* *cells*. Both *WT* and *cAMP-fixed*\* cells were also observed as growing micro-colonies with phase-contrast and fluorescence time-lapse microscopy. The mean growth rates for different *WT* colonies showed an upper range that was similar to the *cAMP-fixed*\* colonies (Fig. S5B). The mean fluorescence intensity per unit area for the CRPr and nCRPr reporters was similar for *WT* and *cAMP-fixed\** cells, on average (Fig. S5C). Overall, these experiments show that *WT* and *cAMP-fixed\** cells display comparable population-mean C-sector stimulation and growth rate, allowing us to study whether the propagation of noise in single cells differs between the *WT* and *cAMP-fixed\** cells.

### The cAMP-CRP system responds to internal stochastic fluctuations

The propagation of and response to cellular noise can be studied by quantifying correlations between fluctuating phenotypic parameters^13,23^. Here we quantify fluctuations in the instantaneous cellular growth rate, λ, by performing phase contrast microscopy at a time resolution of 1 to 1.5 minutes. Fluctuations in CRPr expression levels, *φ*_CRPr_, are quantified by the mVenus fluorescence intensity, using concurrent fluorescence microscopy at intervals ranging from 13.5 to 26 minutes. In *WT* cells, *φ*_CRPr_ fluctuations were negatively correlated with λ fluctuations, as apparent from the negative regression slope in Fig. 1C. In *cAMP-fixed*\* cells, a positive correlation was found instead (Figs. 1D). The difference is statistically significant (p = 0.0031, two sample *t*-test, Fig. S5A). The negative *φ*_CRPr_-λ correlation observed in *WT* cells could reflect the known negative relation between C-sector expression and growth rate under variation of the available carbon sources, also referred to as the C-line^2,4,24^. In line with this hypothesis, this negative correlation is lost in the *cAMP-fixed*\* cells, where the cAMP-CRP feedback is disrupted (Fig. 1D).

To assess whether the *φ*_CRPr_-*λ* correlation changes were due to disruption of CRP regulation, we studied the relationship between *λ* and the expression levels of the reporter not regulated by CRP, *φ*_nCRPr_, as quantified by mCerulean fluorescence intensity. The correlation between *φ*_nCRPr_ and λ was negative for both the *WT* and the *cAMP-fixed*\* cells, and indeed were indistinguishable (Fig. 1E-F, S5A, p = 0.93, Welch’s *t*-test). This similarity is consistent with the above hypothesis, as nCRPr is not regulated by cAMP-CRP in either strain. Thus, the growth correlations of the nCRPr reporter were similar with or without the cAMP-CRP feedback. Conversely, the growth correlations of the CRPr reporter showed a notable shift when this feedback was disrupted, suggesting that the cAMP-CRP system actively responds to internal fluctuations.

### The time-dependent cross-correlations are captured by a mathematical model

To obtain a mechanistic understanding of the observed correlations, we extended the model presented by Kiviet *et al* ^13^. Our aim is not to capture the many known molecular mechanisms of the CRP system, but rather to assess whether the phenomenological relations between variables are sufficient to describe our data. The model is based on linear stochastic differential equations (SDEs) that describe the temporal dynamics of protein production rates (*π*) and concentrations (*φ*), the growth rate (*λ*), and a parameter that reflects the metabolic activity to which the CRP system responds (*M*) (Fig. 2A).

**Figure 2.**
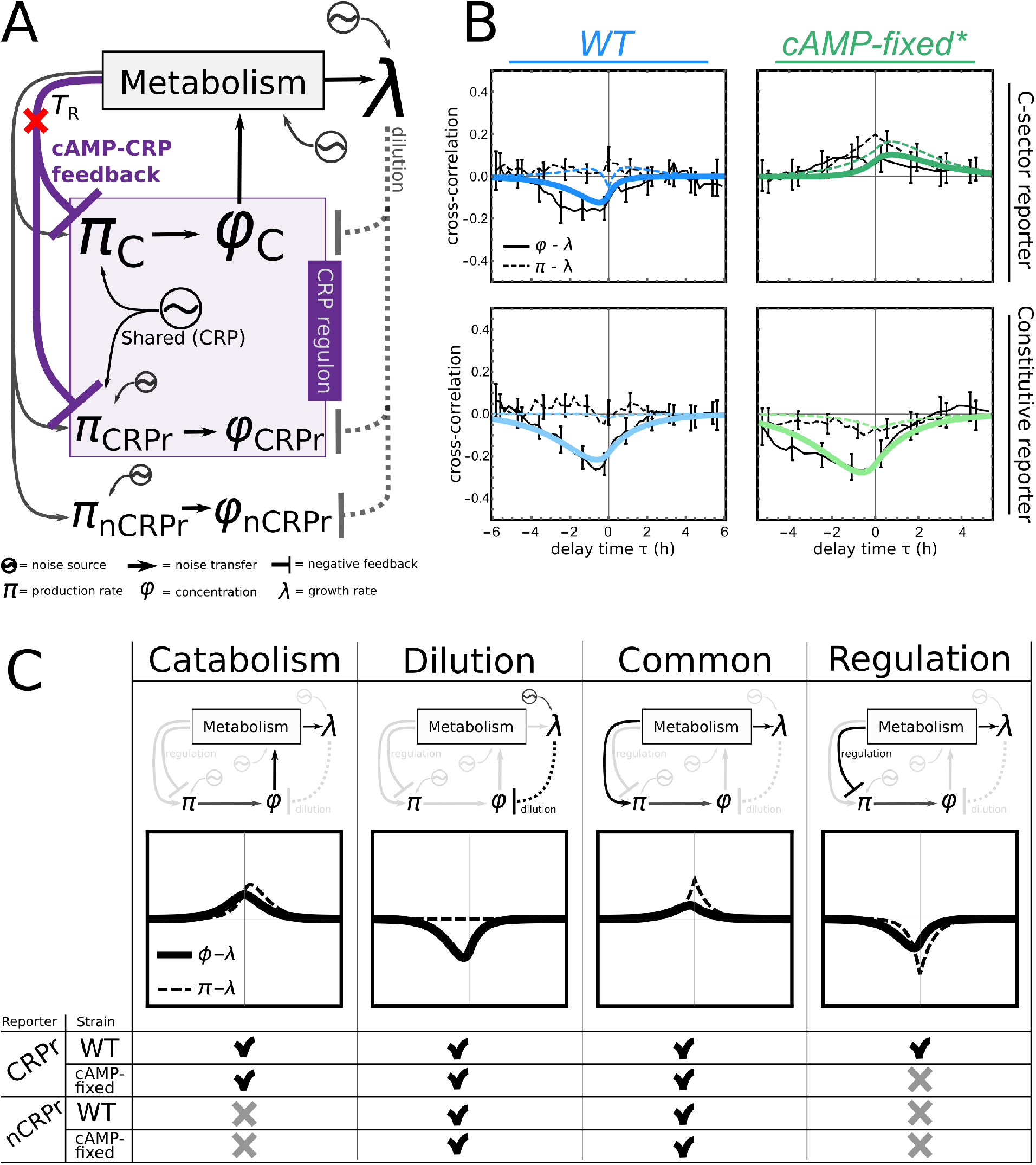
Mathematical model pinpoints dynamical role of regulation. (A) Cartoon of the mathematical model, which considers fluctuations in the growth rate (λ) and the production rates (π) and concentrations (φ) of the C-sector (π_C_ and φ_C_), the C-sector reporter CRPr, and the constitutive reporter nCRPr. Black arrows indicate noise transfer; only fluctuations in φ_C_ affect metabolism. Metabolism affects growth and protein productions rates. Regulation reacts to metabolic fluctuations and transfers to π_C_ and π_CRPr_. In the mutant, regulation is removed (red cross). (B) Cross-correlation functions between the protein production rate π(t) and λ(t) (dashed lines) and between concentrations φ(t) and λ(t), in the wild type and the mutant. Colored lines are model fits, black lines are cross-correlations calculated from data (6 colonies for WT, with n = 3635 cells in total, and 4 colonies for cAMP-fixed*, with n = 6770 cells in total, see Table S2, and Figs. S5 and S7) and error bars indicate standard error (see SI sec 5.2), shown for only some data points. (C) Interpretation and shape of the underlying noise modes that are present in the model. The checks and crosses indicate whether a mode was included in the model’s fit for each experimental condition. The effect of keeping cAMP fixed is reflected by the removal of the regulation mode. Cartoons indicate the direction and route of noise transfer for each specific mode.

We explicitly modeled the expression of the C-sector (*π*_C_ and *φ*_C_), the C-sector reporter (*π*_CRPr_ and *φ*_CRPr_), and the constitutive reporter (*π*_nCRPr_ and *φ*_nCRPr_). Note that the concentrations *φ* depend on the growth rate *λ*, as volume growth dilutes cellular components, while the production rates *π* do not. With this model, we hypothesize that intrinsic stochasticity in *λ, π,* and *M*, as modeled by independent Ornstein-Uhlenbeck noise sources, propagate through the network as defined by the interactions drawn in Fig. 2A, and quantified by coupling coefficients *T*. In particular, it surmises that the cAMP-CRP system propagates internal stochastic fluctuations in constant external conditions, as reflected in the parameter *T_R_*, which directs noise in *M* backwards, to expression of the C-sector genes (Fig. 2A, purple interaction and purple box). This transmission involves fluctuations in metabolite abundance, cAMP synthesis and degradation, and CRP-mediated expression stimulation (Fig. 2A and SI).

The resulting theoretical expressions can be fitted to the experimental data, which also allows one to estimate the transfer coefficients, time scales, and noise amplitudes. Here we focus on fitting the cross-correlation functions *R*, which quantify the correlation between two time series after one is shifted by a delay *τ* and give insights into how noise is transmitted within cellular networks^13,23,25,26^. For example, if noise in signal *A* affects a downstream signal *B* with a fixed time delay, the A-B cross-correlation peaks at a positive *τ*.

For *WT* cells, we found that the *φ*_CRPr_ − *λ* correlation function 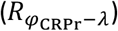 is negative at *τ* = 0 (Fig. 2B), consistent with the negative slope between *φ*_CRPr_ and *λ* (Fig. 1C). For *cAMP-fixed*\* cells, 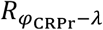 was positive at *τ* = 0 (Fig. 2B), consistent with the positive slope observed in Fig. 1D. A number of other features also became clear. For example, 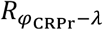 and 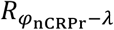 were not only negative in magnitude in *WT* cells, but also peaked at negative delays (*τ* < 0) (Fig. 2B, right). In addition, both 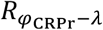 and 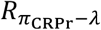 were higher in *cAMP-fixed*\* than in *WT* cells (Fig. 2B, top-right). The cross-correlation functions of the constitutive reporter nCRPr for *WT* and *cAMP-fixed*\* cells were overall very similar (Fig. 2B, bottom).

We simultaneously fitted 8 cross-correlation functions, covering *WT* and *cAMP-fixed*\* cells, the two reporters CRPr and nCRPr, and the production rates *π* and concentrations *φ* (Fig. 2B). To account for the cAMP-CRP feedback disruption, the corresponding transfer parameter *T_R_* was constrained to negative values in *WT* cells but set to zero in *cAMP-fixed*\* cells (Fig. 2A, red cross). All other parameters, including noise transfer parameters, noise amplitudes and timescales were constrained to having the same value for each of the 8 cross-correlation curves (see SI). Despite these strict fitting constraints, the model described the data quantitatively. In particular, it reproduced the shift for the CRPr reporter: from a negative *φ*-*λ* correlation with a negative time delay for *WT* cells, to a nearly flat but slightly positive *φ*-*λ* correlation for *cAMP-fixed*\* cells, and the lack of such a change for the nCRPr reporter (Fig. 2B). Hence, these findings indicate that the cAMP-CRP system actively modulates C-sector expression in response to internal metabolic fluctuations.

### cAMP-CRP noise circulation can be decomposed into distinct noise propagation modes

Next, we further analyzed the model to understand the underlying noise propagation mechanisms. As described above, we postulated coupled (stochastic) differential equations that reflect the stochastic and regulatory dynamics of the CRP-system, and mathematically derived expressions for the cross-correlation functions from this model. Further inspection of these expressions reveal that they are a sum of four noise modes, which we termed the *catabolism, dilution, common*, and *regulation* mode (Fig. 2C and SI). Each mode yields cross-correlation functions of a particular shape and exhibits an amplitude that depends on the amplitudes of the noise sources and transmission parameters; together the modes determine the overall cross-correlation function. The modes describe how emitted noise propagates along particular pathways to two quantities, and hence correlates them (Fig. S2). For instance, in the *catabolism mode*, a stochastic increase in the production rate of a metabolic enzyme leads to higher enzyme concentrations sometime later, and subsequently to a higher growth rate. This mode thus contributes a positive peak at a positive delay time to the *π*-*λ* cross-correlation. In the *dilution mode*, stochastic increases in growth rate leads to increased dilution of all proteins, contributing to the *φ*-*λ* correlation a negative contribution with a negative delay. The *common mode* is the result of fluctuations in general components that directly affect the protein production rate as well as the growth rate. Hence, this mode yields a symmetric *π*-*λ* cross-correlation, while having a negative delay for *φ*-*λ* because of the time needed to change concentrations. Lastly, the *regulation mode* represents noise transferred via the cAMP-CRP system. It shows how stochastic increases in metabolism and growth can transiently limit cAMP-CRP mediated activation of C-sector expression, yielding a symmetric but negative *π*-*λ* cross-correlation, as well as a delayed negative *φ*-*λ* cross-correlation.

The mathematical analysis moreover revealed that not all modes are present in each of the derived cross-correlation functions (Fig 2C, lower table). Interestingly, just the absence and presence of noise modes can already help to qualitatively understand the shape of the experimentally measured cross-correlations for a specific reporter and strain (Fig. 2B-C). First, the analysis of our model suggests that the CRPr reporter cross-correlation contains a catabolism mode, whereas the nCRPr reporter does not. Note that, although neither reporter directly influences the growth rate, the C-sector reporter CRPr can be seen as a proxy for expression of the C-sector, which does influence the growth rate. Therefore, a part of the catabolism mode of the C-sector can be observed in the CRPr cross-correlations, but not in the cross-correlations for the constitutive reporter. Second, the regulation mode is only present in the cross-correlations for the C-sector reporter CRPr in the wild type, because only this reporter is regulated via the cAMP-CRP regulatory network.

We noted that the (cross-)correlations between *φ*_CRPr_ and *λ* (Fig. 1C and Fig. 2B) and between *φ*_nCRPr_ and *λ* (Fig. 1E, 2B) looked similar in WT cells. The mathematical analysis of the noise propagation model, however, indicates that they are composed of different modes (Fig. 2C).For the constitutive reporter nCRPr, in both *WT* and *cAMP-fixed*\* cells, catabolism and regulation modes are absent and the main contribution comes from the dilution mode. Cross-correlations of the C-sector reporter CRPr, on the other hand, additionally contain the catabolism and regulation modes, which largely cancel out, resulting in *WT* correlations with a shape similar to those of the nCRPr reporter. In the *cAMP-fixed*\* cells, the negative regulation mode is absent in CRPr correlations, and the catabolism mode becomes visible, resulting in a positive (cross-)correlation.

Taken together, these observations show that temporal dynamics can be modelled as a linear combination of modes, consistent with the idea that multiple cellular processes, including metabolism and regulation, shape cellular heterogeneity.

### Mechanistic model predicts noise propagation for non-optimal C-sector expression

To further test the model, we sought to describe the effects of changes in the population-mean expression of the C-sector. Hence, we examined cAMP-fixed cells with cAMP concentrations below and above the optimal value, here referred to as *cAMP-fixed*^low^ and *cAMP-fixed*^high^ cells. Consistently, the measured population-mean expression of the C-sector reporter CRPr was below or above that of *cAMP-fixed*\* cells, respectively (Fig. 3A, black dots). Notably, the constitutive nCRPr reporter showed the opposite: the mean expression was higher in *cAMP-fixed*^low^, and lower in *cAMP-fixed*^high^ cells (Fig 3B, red and orange clouds). These observations are consistent with limitations to the size of the overall proteome within cells: when the C-sector becomes larger, the other proteins must decrease in abundance if the total is constrained^2^ (Figs. S10 and 3C). The slow growth of *cAMP-fixed*^low^ cells is consistent with the C-sector becoming growth-limiting when under-expressed, while the slow growth of *cAMP-fixed*^high^ cells is in line with the metabolic costs of superfluously over-expressing the C-sector^5,13,27^.

**Figure 3.**
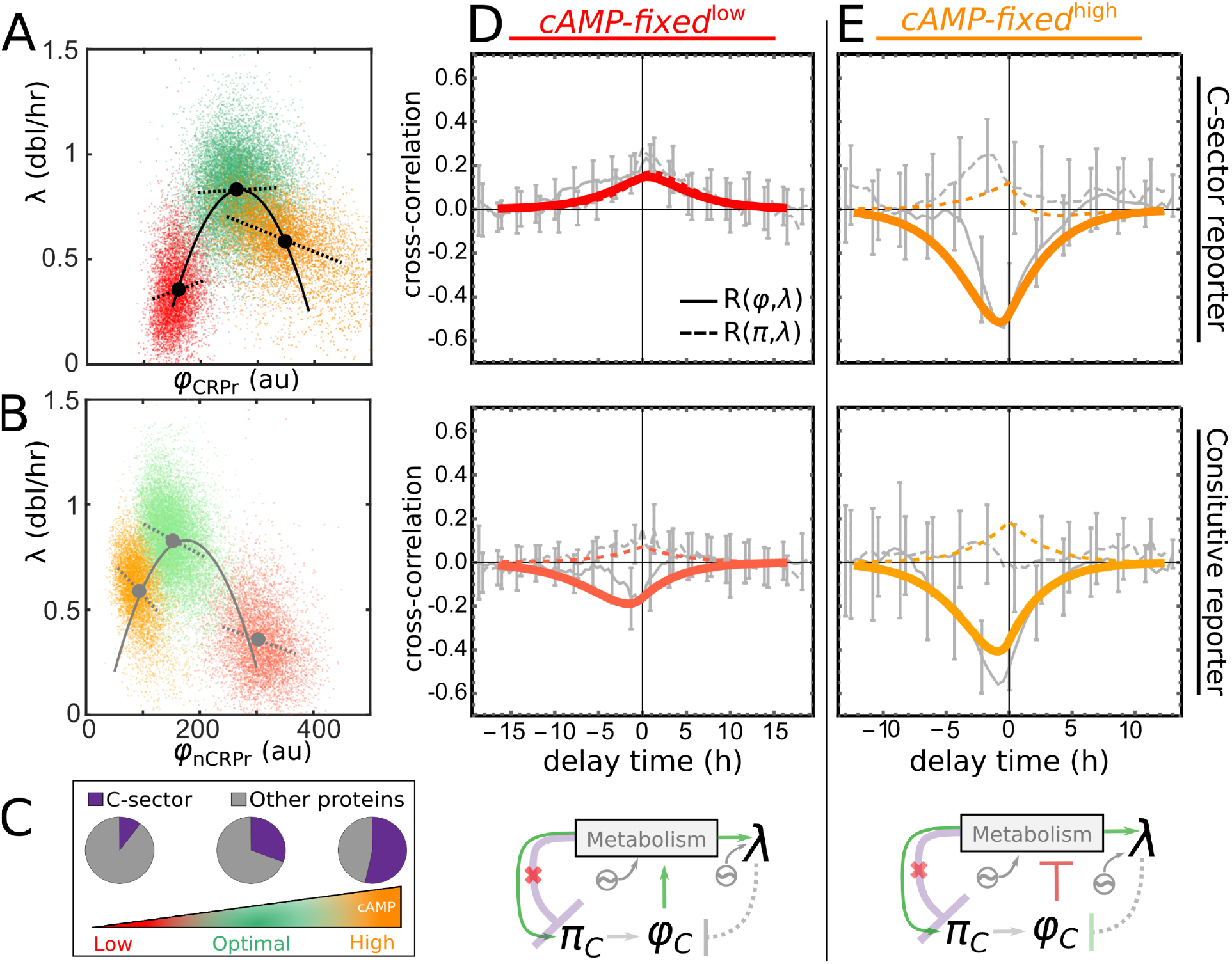
Excessive or insufficient cAMP dampens growth and changes noise-mode amplitudes. (A) Scatter plot of the C-sector reporter against the growth rate, under three conditions: low external cAMP (red cloud), optimal cAMP (green cloud, same condition as in Fig 1D, 2B), high external cAMP (orange cloud). Black dots indicate averages, dashed lines are linear regressions (extending 2 std to each side), black curve is a 2nd order polynomial fit to the means. (B) Same as in B, but then for the constitutive reporter. Grey parabola is calculated from a sum constraint of both reporters (SI sec 7). (C) Cartoon showing how increasing the external cAMP concentration increases the size of the C-sector in the mutant strain, but represses other proteins. (D-E) Measured cross-correlations (grey lines with error bars indicate se) for both reporters for low cAMP (80 μM) and high cAMP (2000 μM), together with model predictions (colored lines) resulting from minimal parametric changes compared to the wild type fit. Model cartoons (bottom panels) indicate changes in transfer parameters (green: increase, red: decrease) with respect to cAMP-fixed* cells (see SI sec 6.2). Data from two microcolonies is shown for low and high cAMP, with respectively n=1788, and n=2274 cells in total, Table S2.

Given these population-mean changes, the model (Fig. 2A) yielded several predictions for the stochastic dynamics. In the *cAMP-fixed*^low^ cells, the growth-limitation of the under-expressed C-sector should increase noise transfer from the C-sector enzyme concentration *φ*_*C*_ to metabolism *M*, and on to the growth rate *λ* and protein production rates *π* (Fig 3D, bottom-left). The associated increases in the transfer coefficients predict overall increases in the *φ*-*λ* and *π*-*λ* correlation functions, owing to increased amplitude of the catabolism mode, while the dilution and common modes remain largely unchanged (Fig. 2C, S6A). We indeed observed that the CRPr correlations, which were positive for *cAMP-fixed*\* cells, had further increased in magnitude, while the nCRPr correlations became less negative (Fig. 3D, S6A). These findings indicate that transient upward fluctuations in C-sector expression can alleviate metabolic bottlenecks caused by mis-regulated C-sector expression, which is on average below the optimum, and hence produce larger increases in growth rate than at optimal C-sector expression.

In the *cAMP-fixed*^high^ cells, the burden of superfluous C-sector expression implies that it now negatively affects metabolism and growth. The corresponding change from positive to negative values for the transfer parameter from *φ*_C_ to M (Fig. 2A) predicts *φ*-*λ* correlations that are strongly negative (Fig. S6B, SI sec 5.2), as now only the weaker common mode yields positive correlations, while both the catabolism and dilution modes are negative (Fig. 2C). The *π*-*λ* correlations are predicted to remain positive however, as *M* couples to *λ* and *π*, which positively correlates them (Fig. 2A). The experiments indeed showed strongly negative *φ*-*λ* correlation (Fig. 3E, solid lines), while *π*-*λ* are positive or negligible (Fig. 3E, dashed lines), in line with these predictions (Fig. S6B). More quantitative fits (Fig. 3D, E) were obtained by increasing the noise amplitudes of the reporters, which decorrelates the signals (Fig. 3D, E, SI sec 5.2). Possibly, such changes in noise amplitudes are caused by changes in average expression levels and mean growth rate, as noise amplitudes tend to increase with the mean^10,28,29^. These experiments indicate that stochastic variations in superfluous expression can cause growth penalties, and that our model captures key aspects of the stochastic dynamics.

The data also showed notable distinctions between population-mean and single-cell behaviors. For instance, regression lines through single-cells clouds (Fig. 3A and B, dashed lines) were typically not tangent to the curves through the population mean values (Fig. 3A and B, solid lines, Fig. S10). Moreover, they were even observed to differ in sign (Fig. 3B, orange cloud). It is also of interest to note that the trade-off between C-sector and other proteins, which we here also observe for the population mean (Fig. 3C), is not obeyed at the single cell level at high cAMP, as cells show stochastic expression increases in the reporters for both the C-sector and the other proteins (Fig. 3A and B, orange clouds).

## Discussion

It has become increasingly clear over the past decade that metabolic networks exhibit stochastic fluctuations^13,15,17,30^. Metabolic networks are also well known to contain numerous regulatory mechanisms that allow cells to react to external changes, which raise the question whether they also act in response to internal noise. Addressing it is critical to understanding whether metabolic homeostasis requires continuous regulatory adjustments, and to elucidating the functional relevance of known regulatory mechanisms. Here, we studied this issue for the cAMP-CRP metabolic regulation system, by experimentally disrupting noise transmission while maintaining the proper population mean activity. Using single cell measurements and mathematical modeling, we found that the cAMP-CRP system modulates the expression of the large group of metabolic enzymes called the C-sector in response to stochastic variations within the metabolic network. Our quantitative approach allowed us to reveal the complex noise circulation pathways within the cAMP-CRP system, which can nonetheless be dissected into distinct and additive noise propagation modes. The latter describe how cAMP-CRP noise regulation relates to catabolic activity, dilution by volume growth, and generic metabolic activity that couples expression and growth in dynamic terms.

This work builds on a growing literature that studies noise in gene expression in the context of metabolism and growth^13,26,31–36^, as well as earlier, theoretical work that mainly focused on metabolic noise propagation within particular regulatory networks^37–39^ and their influence on expression noise. Our modelling framework was chosen to address noise propagation and feedback but is not well-suited to incorporate environmental changes. Modelling approaches that integrate both would help to further understand the differences between responses to external and internal variations. One may first establish the non-linear functions that describe the population-mean responses of a regulatory network to external variations, supported by bulk experiments that dynamically probe expression responses to environmental changes in growth media, antibiotics, or C-sector expression^40,41^. Whether the linearization of such functions properly predict the noise coupling under fixed environmental conditions is unclear. Deviations between the two could originate from differences in the timescales of the variations, how they relate to other fluctuations occurring simultaneously, or the different way in which they tax the overall proteome production capacity of the cell. More generally, there are still many important open questions about the extent to which cells continuously re-adjust their proteome and ribosome abundance in response to stochastic variations in cellular metabolism.

While our phenomenological model reproduced the key experimental findings, it is interesting to speculate about relations to additional mechanisms. The C-sector is subject to global regulation, but each gene within it can be affected by intrinsic noise and other gene-specific variations, which contribute to heterogeneity in metabolite concentrations and metabolic fluxes^17,42,43^. Metabolic noise within our model therefore may be viewed as the compound result of the expression and resulting metabolic noise of many enzymes. Metabolic noise is detected by the cAMP-CRP feedback and transmitted backwards to the entire C-sector that drives metabolism. Hence, noise originating in many pathways can reverberate globally through the cell: multiple cellular processes can transmit, modify and amplify metabolic noise, whatever its provenance, such that it becomes difficult to disentangle source from intermediary.

While CRP is an important master regulator, many other secondary messengers and regulatory mechanisms in metabolic networks are known to respond to external growth conditions. For instance, (p)ppGpp is a crucial global modulator of protein expression, cellular growth, ribosome biogenesis, and cell size upon changes in growth media^44–46^. The notion that metabolic regulation mechanisms can also serve to detect and transmit stochastic fluctuations of metabolites, as we show here, may well apply to these and other regulators. Since stochastic fluctuations could occur in any metabolite, including those that exert allosteric control, our findings suggest that noise may propagate through the cellular networks via diverse and complex feedback mechanisms. We surmise that an understanding of the elementary underlying mechanisms is critical to understanding how cells achieve metabolic homeostasis, as well as how they diversify into heterogenous populations.

## Methods

### Strains

All strains used were based on wild type strain MG1655 (CGSC 8003, bBT12). The CRPr and nCRPr promoters were based on the lac operon promoter, with respectively the lacI binding site or both lacI and CRP binding site scrambled^5^. To obtain the C-sector reporter (CRPr) and constitutive reporter (nCRPr), we fused these promoters to mCerulean and mVenus sequences respectively. The reporters where then inserted into the chromosomes of the bBT12 strain and a cyaA cpda null mutant strain constructed earlier (bBT80), using a lambda red protocol^5^. See Table S3 for strain details and Fig. S11 for promoter sequences.

### Bulk measurements

To determine growth rates of the cyaA cpdA null mutant strain (strain ASC1004, also referred to as cAMP-fixed) at different cAMP concentrations, this strain was inoculated from a freeze mix stock (kept at -80 C) into TY medium, and grown for several hours until exponential growth was achieved. The culture was then diluted (>1000x) into M9 minimal medium supplemented with 0.2 mM uracil, 0.1% lactose, 800 μM cAMP and grown O/N to allow cells to adjust to the lactose medium. Subsequently, the culture was inoculated into separate wells each containing M9 medium with a different final concentration of cAMP (3.1 μM, 8.7 μM, 25.8 μM, 82.7 μM, 272.1 μM, 903.1 μM, 3004.3 μM and 10001.3 μM cAMP, with identical supplements, on a 96 well plate). The samples were then grown for several hours in a Wallac 1420 VICTOR3 Multilabel Counter (Perkin Elmer) to record OD values over time in triplicate. Everything was conducted at 37 C.

### Single cell experiments

Micro-colonies of cells were grown on gel pads, imaged under a microscope, and analyzed by computer as described earlier^13,43^. Briefly, polyacrylamide gel pads (approx. 5 mm x 5 mm x 1 mm in size) were pre-soaked in M9 minimal medium supplemented with lactose (0.01% g/mL), uracil (0.2 mM), Tween20 (0.001%) and the desired concentration of cAMP (Sigma Aldrich) if applicable. Pads were placed in a sealed glass chamber created by a microscope slide and a 2nd glass cavity slide, covered by a glass cover slip. Cells were pre-grown overnight in the same medium, and 1 μl of exponentially growing culture (OD 0.005) was then inoculated on the gel pad at the start of the experiment. Everything was done at 37 °C, and the glass chamber with pad and cells was then placed in a customized scaffold, and imaged under a microscope with a customized incubation chamber at 37 °C. For the *WT, cAMP-fixed*^low^, *cAMP-fixed*\* and *cAMP-fixed*^high^ conditions, we respectively processed time series data from 6, 2, 4 and 2 micro-colonies.

### Microscopy

We used a Nikon, TE2000 microscope, equipped with 100X oil immersion objective (Nikon, Plan Fluor NA 1.3), cooled CMOS camera (Hamamatsu, Orca Flash4.0), xenon lamp with liquid light guide (Sutter, Lambda LS), GFP, mCherry, CFP and YFP filter set (Chroma, 41017, 49008, 49001 and 49003), computer controlled shutters (Sutter, Lambda 10-3 with SmartShutter), automated stage (Märzhäuser, SCAN IM 120 × 100) and an incubation chamber (Solent) allowing precise 37 °C temperature control. An additional 1.5X lens was used, resulting in images with pixel size of 0.0438 μm. The microscope was controlled by MetaMorph software, which allowed us to automatically take pictures at set intervals. Image acquisition intervals were adjusted to doubling times to obtain multiple fluorescent images per cell cycle; phase contrast images were taken every 60-90 seconds, CFP and YFP fluorescent images (150-200 ms exposure time) were taken at intervals ranging from 13.5-26 minutes.

### Image analysis

Series of phase contrast images were analyzed with a custom Matlab (Mathworks) program originally derived from Schnitzcells software ^47^. Cells were segmented and tracked to follow cells and lineages through time. For each frame, cell lengths were determined by fitting a 3rd order polynomial to the curved segmentation regions. Cells were assumed to have a constant width. Growth rates (dbl/hr) were determined by fitting an exponential function to the cell lengths over multiple frames (5 to 9). To determine the production rate per volume, first the sum of the fluorescence signal (a.u.) over all pixels that make up a cell was calculated. If on frame *n* also a fluorescence image was taken, we then calculated the slope of a linear fit through three points *n-l, n*, and *n+l* (where *l* is the frame interval at which fluorescence pictures are taken), the resulting number is divided by the total number of pixels of the cell in frame *n* to obtain the production rate. Concentrations were determined by dividing the sum of the fluorescence signal by the total number of pixels in a cell. To determine scatter plots and correlations, only frames where fluorescence images were taken are considered.

### Cross-correlation analysis

We define the cross-correlation between signals *f* and *g* as 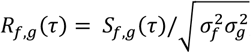, with 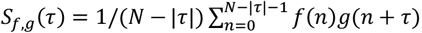, where *n* equals discrete units of time as frame numbers, *τ* is a delay in number of frames, *N* the total number of time points in the data series and 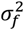 the variance of *f* (equal to *S_f,f_*(0)). See the supplement for more details about data weights and statistics.

### Mathematical model

As mentioned, our model consists of stochastic differential equations, and includes parameters for the protein production rates (*π*), protein concentrations (*ϕ*), metabolism (*M*), and growth rate (*λ*), Ornstein-Uhlenbeck noise sources *N* and noise transfer coefficients *T* that couple equations for *dπ*/*dt*, d*M*/*dt*, and d*λ*/*dt*; concentrations are determined by *dφ*/*dt* = *π* − *φλ*. This model is solved analytically to predict cross-correlations between the quantities. See the supplement for an extensive description of the model, procedures to fit the model to experimental data, a statistical null model for the cross-correlations, and a toy model that describes the mean behavior of *π, ϕ*, and *λ* for CRPr and nCRPr in different conditions as observed in Fig. 3.

### Script and data availability

**Single-cell data,**Matlab scripts and Mathematica notebooks used to create all the figures can be request by the authors, and can be found at: https://github.com/Jintram/DynamicalRegulationBacterialCells.

## References

1. Klumpp, S. & Hwa, T. Bacterial growth: global effects on gene expression, growth feedback and proteome partition. Curr. Opin. Biotechnol. 28, 96–102 (2014).

2. Scott, M., Gunderson, C. W., Mateescu, E. M., Zhang, Z. & Hwa, T. Interdependence of Cell Growth and Gene Expression: Origins and Consequences. Science (80-.). 330, 1099–1102 (2010).

3. Hui, S. et al. Quantitative proteomic analysis reveals a simple strategy of global resource allocation in bacteria. Mol. Syst. Biol. 11, 784 (2015).

4. Schmidt, A. et al. The quantitative and condition-dependent Escherichia coli proteome. Nat. Biotechnol. 34, 104–110 (2016).

5. Towbin, B. D. et al. Optimality and sub-optimality in a bacterial growth law. Nat. Commun. 8, 1–8 (2017).

6. Molenaar, D., van Berlo, R., de Ridder, D. & Teusink, B. Shifts in growth strategies reflect tradeoffs in cellular economics. Mol. Syst. Biol. 5, 323 (2009).

7. Bosdriesz, E., Molenaar, D., Teusink, B. & Bruggeman, F. J. How fast-growing bacteria robustly tune their ribosome concentration to approximate growth-rate maximization. FEBS J. 282, 2029–2044 (2015).

8. Scott, M., Klumpp, S., Mateescu, E. M. & Hwa, T. Emergence of robust growth laws from optimal regulation of ribosome synthesis. Mol. Syst. Biol. (2014).

9. Elowitz, M. B., Levine, A. J., Siggia, E. D. & Swain, P. S. Stochastic Gene Expression in a Single Cell. Science (80-.). 297, 1183–1186 (2002).

10. Taniguchi, Y. et al. Quantifying E. coli proteome and transcriptome with single-molecule sensitivity in single cells. Science 329, 533–538 (2010).

11. Eling, N., Morgan, M. D. & Marioni, J. C. Challenges in measuring and understanding biological noise. Nat. Rev. Genet. 20, 536–548 (2019).

12. Raj, A. & van Oudenaarden, A. Nature, Nurture, or Chance: Stochastic Gene Expression and Its Consequences. Cell 135, 216–226 (2008).

13. Kiviet, D. J. et al. Stochasticity of metabolism and growth at the single-cell level. Nature 514, 376–379 (2014).

14. Taheri-Araghi, S. et al. Cell-size control and homeostasis in bacteria. Curr. Biol. CB 25, 385–391 (2015).

15. Nikolic, N. et al. Cell-to-cell variation and specialization in sugar metabolism in clonal bacterial populations. PLOS Genet. 13, e1007122 (2017).

16. Vasdekis, A. E. & Singh, A. Microbial metabolic noise. WIREs Mech. Dis. 13, e1512 (2021).

17. Fuentes, D. A. F., Manfredi, P., Jenal, U. & Zampieri, M. Pareto optimality between growth-rate and lag-time couples metabolic noise to phenotypic heterogeneity in Escherichia coli. Nat. Commun. 2021 121 12, 1–12 (2021).

18. You, C. et al. Coordination of bacterial proteome with metabolism by cyclic AMP signalling. Nature 500, 301–6 (2013).

19. Kochanowski, K. et al. Few regulatory metabolites coordinate expression of central metabolic genes in Escherichia coli. Mol. Syst. Biol. 13, 903 (2017).

20. Gerosa, L. et al. Pseudo-transition Analysis Identifies the Key Regulators of Dynamic Metabolic Adaptations from Steady-State Data. Cell Syst. 1, 270–282 (2015).

21. Kaplan, S., Bren, A., Dekel, E. & Alon, U. The incoherent feed-forward loop can generate non-monotonic input functions for genes. Mol. Syst. Biol. 4, 203 (2008).

22. Bren, A. et al. Glucose becomes one of the worst carbon sources for E.coli on poor nitrogen sources due to suboptimal levels of cAMP. Sci. Reports 2016 61 6, 1–10 (2016).

23. Dunlop, M. J., Cox, R. S., Levine, J. H., Murray, R. M. & Elowitz, M. B. Regulatory activity revealed by dynamic correlations in gene expression noise. Nat. Genet. 40, 1493–1498 (2008).

24. You, C. et al. Coordination of bacterial proteome with metabolism by cyclic AMP signalling. Nature 500, 301–306 (2013).

25. Thomas, P., Terradot, G., Danos, V. & Weiße, A. Y. Sources, propagation and consequences of stochasticity in cellular growth. Nat. Commun. 9, 1–11 (2018).

26. Kleijn, I. T., Krah, L. H. J. & Hermsen, R. Noise propagation in an integrated model of bacterial gene expression and growth. PLOS Comput. Biol. 14, e1006386 (2018).

27. Dekel, E. & Alon, U. Optimality and evolutionary tuning of the expression level of a protein. Nature 436, 588–592 (2005).

28. Bar-Even, A. et al. Noise in protein expression scales with natural protein abundance. Nat. Genet. 38, 636–643 (2006).

29. Wolf, L., Silander, O. K. & van Nimwegen, E. Expression noise facilitates the evolution of gene regulation. Elife 4, 1–48 (2015).

30. Vasdekis, A. E. et al. Eliciting the impacts of cellular noise on metabolic trade-offs by quantitative mass imaging. Nat. Commun. 2019 101 10, 1–11 (2019).

31. Martino, D. De, Capuani, F. & Martino, A. De. Growth against entropy in bacterial metabolism: the phenotypic trade-off behind empirical growth rate distributions in E.coli. Phys. Biol. 13, 36005 (2016).

32. Susman, L. et al. Individuality and slow dynamics in bacterial growth homeostasis. Proc. Natl. Acad. Sci. 115, E5679--E5687 (2018).

33. Singh, A., Razooky, B. S., Dar, R. D. & Weinberger, L. S. Dynamics of protein noise can distinguish between alternate sources of gene-expression variability. Mol. Syst. Biol. 8, 607 (2012).

34. Lin, J. & Amir, A. From single-cell variability to population growth. Phys. Rev. E 101, 12401 (2020).

35. Shahrezaei, V. & Marguerat, S. Connecting growth with gene expression: of noise and numbers. Curr. Opin. Microbiol. 25, 127–135 (2015).

36. De Martino, D., MC Andersson, A., Bergmiller, T., Guet, C. C. & Tkačik, G. Statistical mechanics for metabolic networks during steady state growth. Nat. Commun. 9, 2988 (2018).

37. Smith, M., Ghusinga, K. R. & Singh, A. Comparison of feedback strategies for noise suppression in protein level. in Proceedings of the American Control Conference vols 2019-July 1513–1518 (2019).

38. Hermsen, R., Erickson, D. W. & Hwa, T. Speed, Sensitivity, and Bistability in Auto-activating Signaling Circuits. PLoS Comput Biol 7, e1002265 (2011).

39. Kittisopikul, M. & Süel, G. M. Biological role of noise encoded in a genetic network motif. Proc. Natl. Acad. Sci. 107, 13300–13305 (2010).

40. Erickson, D. W. et al. A global resource allocation strategy governs growth transition kinetics of Escherichia coli. Nature 551, 119–123 (2017).

41. Korem Kohanim, Y. et al. A Bacterial Growth Law out of Steady State. Cell Rep. 23, 2891–2900 (2018).

42. Tonn, M. K., Thomas, P., Barahona, M. & Oyarzún, D. A. Stochastic modelling reveals mechanisms of metabolic heterogeneity. Commun. Biol. 2, 1–9 (2019).

43. Wehrens, M., Büke, F., Nghe, P. & Tans, S. J. Stochasticity in cellular metabolism and growth: Approaches and consequences. Curr. Opin. Syst. Biol. 8, 131–136 (2018).

44. Büke, F., Grilli, J., Cosentino Lagomarsino, M., Bokinsky, G. & Tans, S. J. ppGpp is a bacterial cell size regulator. Curr. Biol. (2022) doi:10.1016/J.CUB.2021.12.033.

45. Steinchen, W., Zegarra, V. & Bange, G. (p)ppGpp: Magic Modulators of Bacterial Physiology and Metabolism. Front. Microbiol. 0, 2072 (2020).

46. Zhu, M. & Dai, X. Growth suppression by altered (p)ppGpp levels results from non-optimal resource allocation in Escherichia coli. Nucleic Acids Res. 47, 4684–4693 (2019).

47. Young, J. W. et al. Measuring single-cell gene expression dynamics in bacteria using fluorescence time-lapse microscopy. Nat. Protoc. 2011 71 7, 80–88 (2011).

