## Supplementary Information for "The interplay between metabolic stochasticity and regulation in single *E. coli* cells"

August 29, 2022

**6 Toy model of the means of the two reporters** **19**

**7 Supplementary Figures** **21**

*These authors contributed equally.

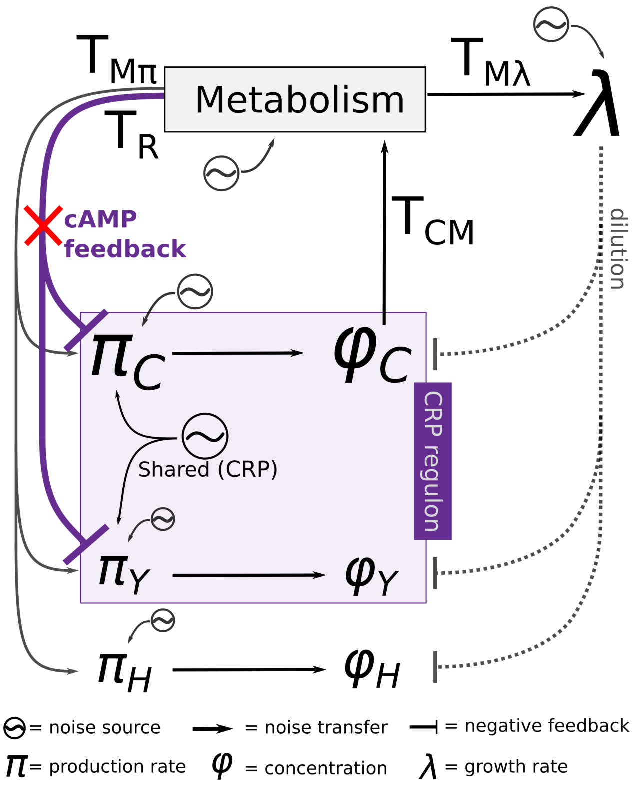

*Figure S1:* Diagram of proposed mathematical model reflecting the biological wiring of a bacterial cell. The model includes the C reporter (Y) and constitutive reporter (H). Intrinsic noise in the protein production rates, metabolism and the growth rate is modeled as independent Ornstein-Uhlenbeck noise sources. An additional shared noise source (here called $N_s$) acts on both $\pi_C$ and $\pi_Y$. $N_s$ represents noise in the sensory mechanism (for example fluctuations of the CRP-concentration). The transfer from Metabolism ($M$) to protein production is the sum of general transfer ($T_{M\pi}$) and regulatory transfer ($T_R$), which only acts on the C-sector and its reporter, Y.

### 1 Mathematical analysis of the noise model

**Note on notation:** Where the main text talks about the C-sector reporter (CRPr) and the not-CRP-regulated constitutive reporter (nCRPr), this document uses the notation 'Y' for the C-sector reporter and 'H' for the constitutive reporter.

#### 1.1 Model definition and parameter interpretation

Similar to Kiviet *et al.* [1] a linear noise propagation model can be constructed using the diagram of Fig. S1. The model captures the interplay between the production rate ($\pi$) of certain proteins, the concentrations ($\phi$) of these proteins, metabolism ($M$) and the growth rate ($\lambda$).

We model the total C-sector in a coarse-grained manner as a single protein, C, whose fluctuations directly influence (the rate of) metabolism. Additionally, we model the two fluorescent reporters, for ease of notation here called Y and H, that are not metabolically active and hence do not control metabolism.

We assume that the variables fluctuate around a particular average state $\{\lambda_0, M_0, \vec{\pi}_0, \vec{\phi}_0\}$; the model describes the dynamics of small, relative deviations of each variable from its respective average, $\frac{\delta X(t)}{X_0} := \frac{X(t)-X_0}{X_0}$. (Below, the explicit time dependence will often be omitted.) These deviations, referred to as "noise", propagate through the system according to the arrows in Fig. S1. For example, fluctuations in the production rate $\pi_C$ affect $\phi_C$, whose fluctuations affect $M$. Via $M$ the noise then further transfers to the production rates and to the growth rate. Noise in the growth rate in turn affects the dilution of all proteins (dashed line). The logarithmic gains/transfer coefficients $T_{\text{AB}}$ describe the strength of noise propagation from $A$ to $B$.

The role of metabolism is crucial in our model because many noise routes pass through metabolism, but also because the cAMP-CRP regulatory network reacts to metabolic fluctuations. The interpretation of $M$ is therefore not straightforward. $M$ could be interpret as the rate of metabolism. However, we assume that with a higher rate of metabolism, the concentrations of certain internal metabolites (in particular keto-acids such as OAA) also increase. The metabolite OAA is known to inhibit the production of cAMP in wild type cells [2] (see Fig. 1A of the main text), therewith triggering the cAMP-CRP regulatory feedback when $M$ increases (represented by the regulatory parameter $T_R$ in the model, Fig. S1). Although we model metabolism with a single variable $M$, it thus represents both the rate of metabolism and the saturation level of metabolic precursors such as OAA.

Certain variables are directly influenced by phenomenological, independent colored noise sources, notated as $N_i$: Ornstein–Uhlenbeck Noise sources that each have a specific timescale. These noise sources offer a way to model the combined effect of many stochastic processes that together influence the dynamics of certain variables. For example, $N_{\pi_Y}$ summarizes the stochastic component of processes that intrinsically influence the production rate of the $Y$ reporter, such as transcription and translation. $N_M$, the noise source acting on metabolism, is primarily the result of fluctuations in the concentration of individual protein species that are part of the C-sector. The fluctuations of all these proteins together are expected to cause the metabolic flux to fluctuate, even if the total size of the C-sector is roughly constant. Fluctuations in the concentrations of individual proteins are diluted via cytoplasm growth, so that the $N_M$ noise source has a timescale close to $\lambda_0$. $N_\lambda$ has a similar interpretation. The noise source $N_s$ equally affects the production of the C-sector and the C-sector reporter Y. Therewith, the noise source $N_s$ is mainly interpreted as fluctuations in the production of CRP-regulated proteins caused by fluctuations in the concentration of CRP itself. Even when the cAMP concentration is experimentally kept fixed in the mutant strain, fluctuations in the CRP concentration are expected to influence the cAMP-CRP concentration and therewith the production rate of both the C-sector and its reporter Y.

Based on the considerations above, we arrive at the following set of equations:

$$\frac{\delta\lambda}{\lambda_0} = N_\lambda + T_{\mathrm{M}\lambda}\frac{\delta M}{M_0}, \tag{1}$$

$$\frac{\delta M}{M_0} = N_{\mathrm{M}} + T_{\mathrm{CM}}\frac{\delta\phi_C}{\phi_{C,0}}, \tag{2}$$

$$\frac{\delta\pi_C}{\pi_{0,C}} = N_{\pi_{\mathrm{C}}} + N_s + (T_{\mathrm{M}\pi} + T_R)\frac{\delta M}{M_0}, \tag{3}$$

$$\frac{\delta\pi_Y}{\pi_{0,Y}} = N_{\pi_{\mathrm{Y}}} + N_s + (T_{\mathrm{M}\pi} + T_R)\frac{\delta M}{M_0}, \tag{4}$$

$$\frac{\delta\pi_H}{\pi_{0,H}} = N_{\pi_{\mathrm{H}}} + T_{\mathrm{M}\pi}\frac{\delta M}{M_0}, \tag{5}$$

$$\left(\frac{\delta\dot{\phi}_C}{\lambda_0\phi_{0,C}}\right) = \frac{\delta\pi_C}{\pi_{0,C}} - \frac{\delta\phi_C}{\phi_{0,C}} - \frac{\delta\lambda}{\lambda_0}, \tag{6}$$

$$\left(\frac{\delta\dot{\phi}_Y}{\lambda_0\phi_{0,Y}}\right) = \frac{\delta\pi_Y}{\pi_{0,Y}} - \frac{\delta\phi_Y}{\phi_{0,Y}} - \frac{\delta\lambda}{\lambda_0}, \tag{7}$$

$$\left(\frac{\delta\dot{\phi}_H}{\lambda_0\phi_{0,H}}\right) = \frac{\delta\pi_H}{\pi_{0,H}} - \frac{\delta\phi_H}{\phi_{0,H}} - \frac{\delta\lambda}{\lambda_0}. \tag{8}$$

The last three equations are liberalizations of the time derivative of the protein concentrations ($\dot{\phi}_i = \pi_i - \lambda\phi_i$), see also [1]. The dynamics of each noise source $N_x$ is given by its stochastic differential equation,

$$\mathrm{d}N_x = -\beta_x N_x \mathrm{dt} + \theta_x \mathrm{dW_t}, \tag{9}$$

where $\beta_x$ is the noise source's timescale, $\theta_x$ its amplitude, and $W(t)$ a Wiener Process.

#### 1.2 Metabolic timescale, solution in Fourier space

The system of equations 1-8 above can be solved in Fourier space. For each variable $A(t)$ we denote the Fourier Transform of $\delta A(t)$ as $\widetilde{A}(\omega)$. Next, we introduce a metabolic rate $\lambda_E$, that scales with the mean growth rate. This rate is related to the time it takes before the protein concentration equilibrates with its protein production rate (note that producing proteins takes time, such that a higher production rate now, only results in a higher concentration some time later). During exponential growth, proteins are generally produced at a (mean) rate proportional to $\lambda_0$, such that production and dilution are balanced. In effect, fluctuations in protein concentrations generally decay on a timescale of $\lambda_0$. However, in the case of C-sector proteins, synthesized proteins directly influence the rate of metabolism (if $T_{CM} \neq 0$), and (depending on the parameters $T_R, T_{M\pi}$ and $T_{M\lambda}$) metabolism in turn influences the production rate and protein concentration. This changes the timescale of C-sector fluctuations from $1/\lambda_0$ to $1/\lambda_E$, where $\lambda_E$ is defined as: $\lambda_E = \lambda_0(1 - T_{CM}T_{MC})$, with $T_{MC} := T_R + T_{M\pi} - T_{M\lambda}$.

Using this notation, we solve the linear system on the previous page in Fourier Space and arrive at:

$$\begin{cases} \frac{\widetilde{\delta\lambda}}{\lambda_0} &= N_\lambda + T_{M\lambda}N_M + \frac{\lambda_0}{\lambda_E + i\omega}\left(T_{CM}T_{M\lambda}\right)\left[N_{\pi_C} + N_s + T_{MC}N_M - N_\lambda\right], \\ \frac{\widetilde{\delta\phi_C}}{\phi_{C0}} &= \frac{\lambda_0}{\lambda_E + i\omega}\left[N_{\pi_C} + N_s + T_{MC}N_M - N_\lambda\right], \\ \frac{\widetilde{\delta\pi_C}}{\pi_{C0}} &= N_{\pi_C} + N_s + (T_R + T_{M\pi})N_M + \frac{\lambda_0}{\lambda_E + i\omega}T_{CM}(T_R + T_{M\pi})\left[N_{\pi_C} + N_s + T_{MC}N_M - N_\lambda\right]. \end{cases} \tag{10}$$

$$\begin{cases} \frac{\widetilde{\delta\phi_Y}}{\phi_{y,0}} &= \frac{\lambda_0}{\lambda_0+i\omega}\left[N_{\pi_Y}+N_s+T_{MC}N_M-N_\lambda\right]+\frac{\lambda_0^2T_{CM}T_{MC}}{(\lambda_E+i\omega)(\lambda_0+i\omega)}\left[N_{\pi_C}+N_s+T_{MC}N_M-N_\lambda\right], \\ \frac{\widetilde{\delta\pi_Y}}{\pi_{y,0}} &= N_{\pi_Y}+N_s+(T_R+T_{M\pi})N_M+\frac{\lambda_0}{\lambda_E+i\omega}T_{CM}\left(T_R+T_{M\pi}\right)\left[N_{\pi_C}+N_s+T_{MC}N_M-N_\lambda\right]. \end{cases} \tag{11}$$

$$\begin{cases} \frac{\widetilde{\delta\phi_H}}{\phi_{H,0}} &= \frac{\lambda_0}{\lambda_0+i\omega}\left[N_{\pi_H}+(T_{M\pi}-T_{M\lambda})N_M-N_\lambda\right]+\frac{\lambda_0^2T_{CM}(T_{M\pi}-T_{M\lambda})}{(\lambda_E+i\omega)(\lambda_0+i\omega)}\left[N_{\pi_C}+N_s+T_{MC}N_M-N_\lambda\right], \\ \frac{\widetilde{\delta\pi_H}}{\pi_{H,0}} &= N_{\pi_H}+T_{M\pi}N_M+\frac{\lambda_0}{\lambda_E+i\omega}T_{CM}T_{M\pi}\left[N_{\pi_C}+N_s+T_{MC}N_M-N_\lambda\right]. \end{cases} \tag{12}$$

#### 1.3 Calculated variances and cross-covariances

##### 1.3.1 Calculation of cross-covariance

The solution in Fourier Space (Eqs. 10 - 12) can be used to calculate variances and cross-covariances of the variables. Using the convolution theorem, cross-covariance of variables $A$ and $B$, with time-averages $A_0$ and $B_0$, can be calculated as:

$$\mathcal{X}_{(A,B)}(\tau)=\frac{1}{2\pi}\mathcal{F}^{-1}\left(\frac{\widetilde{A(t-\tau)}^*}{A_0}\frac{\widetilde{B(t)}}{B_0}\right), \tag{13}$$

where $\mathcal{F}^{-1}(.)$ denotes the Inverse Fourier Transform (* here denotes complex conjugation). In the product $\frac{\tilde{A}^*}{A_0}\frac{\tilde{B}}{B_0}$ we can safely ignore terms of $N_iN_j^*$ when $i\neq j$, for they will not contribute to the cross-correlation due to independence of the noise sources. Consequently, the expansion of the product is always linear in the absolute values of the noise sources $|N_i|^2$. This feature is important, because it assures that the cross-covariance can be written as $\chi_{(A,B)}(\tau)=\frac{1}{2\pi}\mathcal{F}^{-1}\left(\sum_i f_i(\omega)|N_i|^2\right)$, where $f_i(\omega)$ are complex functions to be determined and the summation runs over all noise sources. Because the Inverse Fourier Transform is a linear operator, we only need to calculate the Inverse Fourier Transform of each term, $f_i(\omega)|N_i|^2$, separately. The Inverse Fourier Transform of each term is given in section 1.3.4.

##### 1.3.2 Analytical expressions for the coefficients of variation

From the above equations (10 - 13) one can also derive the coefficient of variation of all variables. Concretely, the coefficient of variation of variable $A$ is related to its auto-covariance at zero delay:

$$\eta_A^2:=\sigma_A^2/A_0^2=\mathcal{X}_{(A,A)}(0). \tag{14}$$

For example, the coefficients of variation for $\phi_C$ and $\lambda$ can be calculated using the Inverse Fourier transform as follows:

$$\eta_{\phi_C}^2:=\mathcal{F}^{-1}_{(\tau=0)}\left[\frac{\widetilde{\delta\phi_C}^*}{\phi_{0,C}}\frac{\widetilde{\delta\phi_C}}{\phi_{0,C}}\right]=\mathcal{F}^{-1}_{(0)}\left[\frac{\lambda_0^2}{\lambda_E^2+\omega^2}\left(|N_{\pi_C}|^2+|N_s|^2+T_{MC}^2|N_M|^2+|N_\lambda|^2\right)\right]. \tag{15}$$

$$\eta_\lambda^2=\mathcal{F}^{-1}_{(\tau=0)}\left[|N_\lambda|^2+T_{M\lambda}^2|N_M|^2+\frac{2\lambda_0\lambda_E}{\lambda_E^2+\omega^2}T_{CM}T_{M\lambda}\left(T_{M\lambda}T_{MC}|N_M|^2-|N_\lambda|^2\right)\right] \tag{16}$$

$$+(T_{CM}T_{M\lambda})^2\eta_{\phi_C}^2. \tag{17}$$

(Here we have used the identity $\frac{\lambda_0}{\lambda_E+i\omega} + \frac{\lambda_0}{\lambda_E-i\omega} = \frac{2\lambda_0\lambda_E}{\lambda_E^2+\omega^2}$.) To calculate these (Inverse) Fourier Transforms we use a set of identities that can be found later in this document (Section 1.3.4, Eqs. 28-38). Here we present the results (using the notation $\beta_X^j := \beta_X + \lambda_j$, with $j \in \{0, E\}$):

$$\begin{cases} \eta_{\phi_C}^2 &= \frac{\lambda_0^2}{2\lambda_E}\left(\frac{\theta_{\pi_C}^2}{\beta_{\pi_C}\beta_{\pi_C}^E} + \frac{\theta_s^2}{\beta_s\beta_s^E} + T_{MC}^2\frac{\theta_M^2}{\beta_M\beta_M^E} + \frac{\theta_\lambda^2}{\beta_\lambda\beta_\lambda^E}\right), \\ \eta_\lambda^2 &= \frac{\theta_\lambda^2}{2\beta_\lambda}\left(1 - \frac{2\lambda_0}{\beta_\lambda^E}T_{CM}T_{M\lambda}\right) + \frac{\theta_M^2}{2\beta_M}T_{M\lambda}^2\left(1 + \frac{2\lambda_0}{\beta_M^E}T_{CM}T_{MC}\right) + (T_{CM}T_{M\lambda})^2\eta_{\phi_C}^2, \\ \eta_{\pi_C}^2 &= \frac{\theta_{\pi_C}^2}{2\beta_{\pi_C}}\left(1 + \frac{2\lambda_0}{\beta_{\pi_C}^E}T_{CM}(T_R + T_{M\pi})\right) + \frac{\theta_s^2}{2\beta_s}\left(1 + \frac{2\lambda_0}{\beta_s^E}T_{CM}(T_R + T_{M\pi})\right) \\ &\quad + \frac{\theta_M^2}{2\beta_M}(T_R + T_{M\pi})^2\left(1 + \frac{2\lambda_0}{\beta_M^E}T_{MC}T_{CM}\right) + T_{CM}^2(T_R + T_{M\pi})^2\eta_{\phi_C}^2. \end{cases} \tag{18}$$

$$\begin{cases} \eta_{\phi_Y}^2 &= \frac{\theta_{\pi_Y}^2\lambda_0}{2\beta_{\pi_Y}\beta_{\pi_Y}^0} + \frac{\theta_s^2\lambda_0}{2\beta_s\beta_s^0}\left(1 + T_{CM}T_{MC}\frac{\lambda_0(\beta_s+\lambda_0+\lambda_E)}{\lambda_E\beta_s^E}\right) + \frac{\theta_M^2\lambda_0}{2\beta_M\beta_M^0}T_{MC}^2\left(1 + T_{CM}T_{MC}\frac{\lambda_0(\beta_M+\lambda_0+\lambda_E)}{\lambda_E\beta_M^E}\right) \\ &\quad + \frac{\theta_\lambda^2\lambda_0}{2\beta_\lambda\beta_\lambda^0}\left(1 + T_{CM}T_{MC}\frac{\lambda_0(\beta_\lambda+\lambda_0+\lambda_E)}{\lambda_E\beta_\lambda^E}\right) + \frac{\theta_{\pi_C}^2\lambda_0^2}{2\beta_{\pi_C}\beta_{\pi_C}^0}T_{CM}^2T_{MC}^2\frac{\lambda_0(\beta_{\pi_C}+\lambda_0+\lambda_E)}{\beta_{\pi_C}^E\lambda_E(\lambda_0+\lambda_E)}, \\ \eta_{\pi_Y}^2 &= \frac{\theta_{\pi_Y}^2}{2\beta_{\pi_Y}} + \frac{\theta_s^2}{2\beta_s}\left(1 + 2\frac{\lambda_0}{\beta_s^E}T_{CM}(T_R + T_{M\pi})\right) + \frac{\theta_M^2}{2\beta_M}(T_R + T_{M\pi})^2\left(1 + 2\frac{\lambda_0}{\beta_M^E}T_{CM}T_{MC}\right) \\ &\quad + T_{CM}^2(T_R + T_{M\pi})^2\eta_{\phi_C}^2. \end{cases} \tag{19}$$

$$\begin{cases} \eta_{\phi_H}^2 &= \frac{\theta_{\pi_H}^2\lambda_0}{2\beta_{\pi_H}\beta_{\pi_H}^0} + \frac{\lambda_0^3T_{CM}^2(T_{M\pi}-T_{M\lambda})^2}{2\lambda_E(\lambda_0+\lambda_E)}\left(\frac{\theta_{\pi_C}^2(\beta_{\pi_c}+\lambda_0+\lambda_E)}{\beta_{\pi_c}\beta_{\pi_c}^0\beta_{\pi_c}^E} + \frac{\theta_s^2(\beta_s+\lambda_0+\lambda_E)}{\beta_s\beta_s^0\beta_s^E}\right) \\ &\quad + \frac{\theta_M^2\lambda_0(T_{M\pi}-T_{M\lambda})^2}{2\beta_M\beta_M^0}\left(1 + T_{CM}T_{MC}\frac{\lambda_0(\beta_M+\lambda_0+\lambda_E)}{\lambda_E\beta_M^E}\right) \\ &\quad + \frac{\theta_\lambda^2\lambda_0}{2\beta_\lambda\beta_\lambda^0}\left(1 + T_{CM}(T_{M\pi} - T_{M\lambda})\frac{\lambda_0(\beta_\lambda+\lambda_0+\lambda_E)}{\beta_\lambda^E(\lambda_0+\lambda_E)}\left(T_{CM}(T_{M\pi} - T_{M\lambda})\frac{\lambda_0}{\lambda_E} + 2\right)\right) \\ \eta_{\pi_H}^2 &= \frac{\theta_{\pi_H}^2}{2\beta_{\pi_H}} + \frac{\theta_M^2}{2\beta_M}T_{M\pi}^2\left(1 + 2\frac{\lambda_0}{\beta_M^E}T_{CM}T_{MC}\right) + T_{CM}^2T_{M\pi}^2\eta_{\phi_C}^2. \end{cases} \tag{20}$$

##### 1.3.3 Analytical expressions for the cross-covariances

The cross-covariances (Eq. 13) are best expressed in terms of the functions $A_x^j, B_x, S_x$, and $D_x^{[k]}$ (where $x$ indicates the noise source, $j \in \{0, E\}$ and $k \in \{1, 2, 3\}$). The definition of these functions and some intuitive explanation for $D_x^{[k]}$ is given in SI, sections 1.3.4 and 2.5. Here, we however already use the relation $A_x(t) + A_x(-t) = 2\frac{\lambda_E}{\lambda_0}S_x(t)$).

$$\chi_{(\phi_C,\lambda)}(t) = T_{CM}T_{M\lambda}\left[S_{\pi_C}(t) + S_s(t) + T_{MC}^2S_M(t) + S_\lambda(t)\right] + T_{MC}T_{M\lambda}A_M(t) - A_\lambda(t). \tag{21}$$

$$\begin{aligned} \chi_{(\pi_C,\lambda)}(t) = T_{CM}^2T_{M\lambda}(T_R + T_{M\pi})&\left[S_{\pi_c}(t) + S_s(t) + T_{MC}^2S_M(t) + S_\lambda(t)\right] + (T_R + T_{M\pi})T_{M\lambda}B_M(t) \\ &+ T_{CM}T_{M\lambda}\left[A_{\pi_C}(-t) + A_s(-t)\right] + \frac{2\lambda_E}{\lambda_0}T_{CM}T_{M\lambda}T_{MC}(T_R + T_{M\pi})S_M(t) \\ &- T_{CM}(T_R + T_{M\pi})A_\lambda(t). \end{aligned} \tag{22}$$

$$\chi_{(\phi_Y,\lambda)}(t) = T_{MC}T_{M\lambda}A^0_M(t) - A^0_\lambda(t)$$
$$+ T_{CM}T_{M\lambda}\Big[D^{[1]}_s(t) + T^2_{MC}D^{[1]}_M(t) + D^{[1]}_\lambda(t)\Big] + T_{CM}T_{MC}\Big[T_{MC}T_{M\lambda}D^{[2]}_M(t) - D^{[2]}_\lambda(t)\Big]$$
$$+ T^2_{CM}T_{MC}T_{M\lambda}\Big[D^{[3]}_{\pi_C}(t) + D^{[3]}_s(t) + T^2_{MC}D^{[3]}_M(t) + D^{[3]}_\lambda(t)\Big]. \quad (23)$$

$$\chi_{(\pi_Y,\lambda)}(t) = T_{M\lambda}(T_R + T_{M\pi})B_M(t) + \frac{2\lambda_E}{\lambda_0}T_{CM}T_{MC}T_{M\lambda}(T_R + T_{M\pi})S_M(t)$$
$$+ \quad T_{CM}\Big[T_{M\lambda}A^E_s(-t) - (T_R + T_{M\pi})A^E_\lambda(t)\Big]$$
$$+ \quad T^2_{CM}(T_R + T_{M\pi})T_{M\lambda}\Big[S_{\pi_C}(t) + S_s(t) + T^2_{MC}S_M(t) + S_\lambda(t)\Big]. \quad (24)$$

$$\chi_{(\phi_H,\lambda)} = (T_{M\pi} - T_{M\lambda})T_{M\lambda}A^0_M(t) - A^0_\lambda(t) + T_{CM}T_{M\lambda}\Big[(T_{M\pi} - T_{M\lambda})T_{MC}D^{[1]}_M(t) + D^{[1]}_\lambda(t)\Big]$$
$$+ T_{CM}(T_{M\pi} - T_{M\lambda})\Big[T_{MC}T_{M\lambda}D^{[2]}_M(t) - D^{[2]}_\lambda(t)\Big]$$
$$+ T^2_{CM}(T_{M\pi} - T_{M\lambda})T_{M\lambda}\Big[D^{[3]}_{\pi_C}(t) + D^{[3]}_s(t) + T^2_{MC}D^{[3]}_M(t) + D^{[3]}_\lambda(t)\Big]. \quad (25)$$

$$\chi_{(\pi_H,\lambda)} = T_{M\pi}T_{M\lambda}B_M(t) + \frac{2\lambda_E}{\lambda_0}T_{M\pi}(T_{MC}T_{CM}T_{M\lambda})S_M(t) - T_{CM}T_{M\pi}A_\lambda(-t)$$
$$+ T^2_{CM}T_{M\pi}T_{M\lambda}\Big[S_{\pi_C}(t) + S_s(t) + T^2_{MC}S_M(t) + S_\lambda\Big]. \quad (26)$$

The variances and cross-covariances presented here can be further studied and checked with the Mathematica notebook/supplementary file 'VarianceChecker.nb' (Mathematica 13), which is available online: https://github.com/Jintram/DynamicalRegulationBacterialCells.

##### 1.3.4 Functional forms of the building blocks

In the above equations, $A^j_x, B_x, S_x$ and $D^{[k]}_x$ ($j \in \{0, E\}$ and $k \in \{1, 2, 3\}$) are functional forms that describe the shape of the contribution of a noise source $N_x$ to the cross-covariance. This contribution depends on the noise source, but also on the route via which noise is transferred through the system (Fig. S1). These functions therefore depend on $\theta_x$ and $\beta_x$, the amplitude and timescale of $N_x$ (for clarity of reading we have omitted all the $x$-subscripts below). Additionally, the functions depend on the specific route via which is transferred through the system (Figs. S1 and S2). The functional forms are found by complex integration using Cauchy's Residue Theorem and are given by:

$$A^j(t) = \int \frac{e^{i\omega t}}{2\pi}\frac{\lambda_0}{\lambda_j - i\omega}\frac{\theta^2}{\beta^2+\omega^2}d\omega, \quad \text{for } j \in \{0, E\}, \text{but superscript "E" is implicit} \tag{27}$$

$$= \frac{\theta^2\lambda_0}{2\beta}\begin{cases} \dfrac{2\beta e^{\lambda_j t}}{\beta^2 - \lambda_j^2} - \dfrac{e^{\beta t}}{\beta - \lambda_j}, & \text{for } t < 0, \\ \dfrac{e^{-\beta t}}{\beta_x + \lambda_j}, & \text{for } t \geq 0. \end{cases} \tag{28}$$

$$S(t) = \int \frac{e^{i\omega t}}{2\pi}\frac{\lambda_0^2}{\lambda_E^2 + \omega^2}\frac{\theta^2}{\beta^2+\omega^2}d\omega \tag{29}$$

$$= \frac{\theta^2\lambda_0^2}{2(\beta^2 - \lambda_E^2)}\left(\frac{e^{-\lambda_E|t|}}{\lambda_E} - \frac{e^{-\beta|t|}}{\beta}\right) \tag{30}$$

$$D^{[1]}(t) = \int \frac{e^{i\omega t}}{2\pi}\frac{\lambda_0^2}{(\lambda_E + i\omega)(\lambda_0 - i\omega)}\frac{\theta^2}{\beta^2+\omega^2}d\omega \tag{31}$$

$$= \theta^2\lambda_0^2\begin{cases} \dfrac{e^{\lambda_0 t}}{(\beta^2 - \lambda_0^2)(\lambda_0 + \lambda_E)} - \dfrac{e^{\beta t}}{2\beta\beta_x^E(\beta - \lambda_0)}, & \text{for } t < 0, \\ \dfrac{e^{-\lambda_E t}}{(\beta^2 - \lambda_E^2)(\lambda_0 + \lambda_E)} - \dfrac{e^{-\beta t}}{2\beta\beta^0(\beta - \lambda_E)}, & \text{for } t \geq 0 \end{cases} \tag{32}$$

$$D^{[2]}(t) = \int \frac{e^{i\omega t}}{2\pi}\frac{\lambda_0^2}{(\lambda_E - i\omega)(\lambda_0 - i\omega)}\frac{\theta^2}{\beta^2+\omega^2}d\omega \tag{33}$$

$$= \theta^2\lambda_0^2\begin{cases} \dfrac{e^{\beta t}}{2\beta(\beta - \lambda_0)(\beta - \lambda_E)} + \dfrac{1}{\lambda_0 - \lambda_E}\left(\dfrac{e^{\lambda_E t}}{\beta^2 - \lambda_E^2} - \dfrac{e^{\lambda_0 t}}{\beta^2 - \lambda_0^2}\right), & \text{for } t < 0, \\ \dfrac{e^{-\beta t}}{2\beta\beta^E\beta^0}, & \text{for } t \geq 0 \end{cases} \tag{34}$$

$$D^{[3]}(t) = \int \frac{e^{i\omega t}}{2\pi}\frac{\lambda_0^3}{(\lambda_E^2 + \omega^2)(\lambda_0 - i\omega)}\frac{\theta^2}{\beta^2+\omega^2}d\omega \tag{35}$$

$$= \frac{\theta^2\lambda_0^3}{2}\begin{cases} \left(\dfrac{e^{\beta t}}{\beta(\beta - \lambda_0)(\beta^2 - \lambda_E^2)} - \dfrac{2e^{\lambda_0 t}}{(\beta^2 - \lambda_0^2)(\lambda_0^2 - \lambda_E^2)} + \dfrac{e^{\lambda_E t}}{\lambda_E(\beta^2 - \lambda_E^2)(\lambda_0 - \lambda_E)}\right), & \text{for } t < 0, \\ \dfrac{e^{-\lambda_E t}}{\lambda_E(\lambda_0 + \lambda_E)(\beta^2 - \lambda_E^2)} - \dfrac{e^{-\beta t}}{\beta\beta^0(\beta^2 - \lambda_E^2)}, & \text{for } t \geq 0 \end{cases} \tag{36}$$

$$B(t) = \int \frac{e^{i\omega t}}{2\pi}\frac{\theta^2}{\beta^2+\omega^2}d\omega \tag{37}$$

$$= \frac{\theta^2}{2\beta}e^{-\beta|t|}. \tag{38}$$

In section 2.5 we will further discuss the interpretation of the these functions.

### 2 Analyzing the expressions for the cross-covariance functions

The cross-covariance between two signals $A$ and $B$ contains temporal information about the dynamical interplay of the two signals. For example, if noise first reaches signal $A$ and only later arrives at $B$, their cross-covariance peaks at a positive delay time. Generally, the complicated expressions for the cross-covariance between $A$ and $B$ indeed consists of several terms that each correspond to a different (biophysical) mechanism by which noise can propagate from some noise source to $A$ and $B$. Insight is gained by splitting the mathematical description of the cross-covariance into different 'modes' that can each be interpreted as a different ways in which noise can propagate through the cell. However, such a decomposition is not unique and hence subject to convention. Therefore, we first discuss a decomposition from Kiviet *et al.* [1] using the $\phi_C$-$\lambda$ cross-covariance, $\chi_{\phi_C,\lambda}$ as an example. Next, we propose a new decomposition that is better suited to the system investigated in this work.

#### 2.1 The $\phi_C$-$\lambda$ cross-covariance: Old Decomposition

First, we split up the $\phi_C$-$\lambda$ cross-covariance (see Eq. 21) in the same way as in Kiviet *et al.*, using three modes:

- **Catabolism mode**. This mode contains the contributions of all paths that transfer via catabolism (*i.e.* from $\phi_C$ to $M$ and from $M$ to $\lambda$), where one route ends at the growth rate, and the other at protein concentration:

$$T_{CM}T_{M\lambda}\left(S_{\pi_C} + S_s + T_{MC}^2 S_M + (-1)^2 S_\lambda\right).$$

- **Common mode**. This mode originates from a phenomenological noise source that directly influences both the growth rate and the production rates (which in turn influences the concentrations):

$$T_{MC}T_{M\lambda}A_M.$$

- **Dilution mode**. This mode represents direct transfer from growth to protein concentration:

$$-A_\lambda.$$

#### 2.2 The $C$-$\lambda$ cross-covariance: New decomposition

In the current work, we argue to define the modes slightly different. The major difference is in the interpretation of the *common* mode, which is now not defined as contributions to the cross-correlations originating from a particular noise source, but rather as all contributions that are sensed commonly by all proteins in the cell. Specifically, noise in the production of particular proteins can contribute to fluctuations of $M$, from where it transfers to all proteins commonly. In other words, noise that originates in a particular part of the cell can propagate through the cell and partly *become* common noise, transferring further to all proteins equally. Additionally we introduce the regulation mode, which contains all contributions to the cross-correlations that rely on the regulation and hence scale linearly with $T_R$.

To indicate paths through the cell along which noise transfers, and to therewith know via which biophysical mechanism the noise transfers, we have added red parentheses to the expression of $\chi_{(\phi_C,\lambda)}$. These red parentheses divide the transfer parameters into groups of transfer parameters that together describe a particular path through the cell (see also Figs. S1 and S2).

From every noise source one can indeed follow two paths, one to each of the variables for which the cross-correlation is calculated. (Here we used that the transfer parameter from growth to protein concentrations is −1.) With these conventions, the decomposition of the cross-correlation between $\phi_C$ and $\lambda$ becomes as follows:

- $\chi_{(\phi_C,\lambda)}$:
  1. **Catabolism**: $(T_{CM}T_{M\lambda})\left(S_{\pi_C}+S_s\right)$.
  2. **Dilution**: $(-1)\,A_\lambda$.
  3. **Common**: $(T_{M\pi})\,(T_{M\lambda})\,A_M+(-T_{M\lambda})\,(T_{M\lambda})\,A_M+(T_{MC})\,(T_{MC}T_{CM}T_{M\lambda})\,S_M+(-1)\,(-T_{CM}T_{M\lambda})\,S_\lambda$.
  4. **Regulation**: $(T_R)\,(T_{M\lambda})\,A_M$.

- $\chi_{(\pi_C,\lambda)}$:
  1. **Catabolism**: $T_{CM}T_{M\lambda}\left(A_{\pi_C}(-t)+A_s(-t)\right)$.
  2. **Dilution**: 0.
  3. **Common**: $T_{M\pi}T_{M\lambda}B_M+T_{CM}T_{M\pi}T_{CM}T_{M\lambda}\left[(S_{\pi_C}+S_s+T^2_{MC}S_M+(-1)^2S_\lambda\right]$ $+2\frac{\lambda_E}{\lambda_0}T_{MC}T_{CM}T_{M\pi}T_{M\lambda}S_M-T_{CM}T_{M\pi}A_\lambda$.
  4. **Regulation**: $T_RT_{M\lambda}B_M+T_{CM}T_RT_{CM}T_{M\lambda}\left[S_{\pi_C}+S_s+T^2_{MC}S_M+S_\lambda\right]$ $+2\frac{\lambda_E}{\lambda_0}T_{MC}T_{CM}T_RT_{M\lambda}S_M-T_{CM}T_RA_\lambda$.

The above equations for $\chi_{(\phi_C,\lambda)}$ and $\chi_{(\pi_C,\lambda)}$ apply to both wild type *E. coli* cells and the cAMP-fixed *cyaA cpdA* null mutant studied in the main text. This allows us to pinpoint the role of regulation in shaping the cross-correlation. The parameter $T_R$ that represents the cAMP-CRP regulation feedback is expected to be negative in the wild type ($T_R<0$ due to cAMP-CRP regulation). In the cAMP-fixed strain however, the negative feedback is abolished and $T_R=0$. Such a parametric switch will qualitatively change the cross-correlations $R_{(\phi_C,\lambda)}$ and $R_{(\pi_C,\lambda)}$. For example, a negative contribution with functional form of $A_M$ (asymmetrically, left-skewed function with timescale $\lambda_0$) is effectively removed from $R_{(\phi_C,\lambda)}$.

Note that $T_R$ however not only influences the presence and absence of modes. Additionally, $T_R$ also influences the coefficients of variation (Eqs. 18 - 20), and amplitudes and timescales of the other modes via $\lambda_E:=\lambda_0(1-(T_R+T_{M\pi}-T_{M\lambda}))$. Most modes are therefore expected to show slight, quantitative differences between wild type and cAMP-fixed cells, but only the regulation mode will show a strong qualitative difference: it is completely absent in cAMP-fixed cells.

#### 2.3 Summary of the new decomposition for the reporters

##### 2.3.1 $\chi(\phi_Y,\lambda)$

1. **Catabolism**: $T_{CM}T_{M\lambda}D^{[1]}_s$.
2. **Dilution**: $-A^0_\lambda$.
3. **Common**: $(T_{M\pi}-T_{M\lambda})T_{M\lambda}A^0_M+T_{CM}T_{MC}\left(T_{MC}T_{M\lambda}D^{[2]}_M-D^{[2]}_\lambda\right)+T_{CM}(T_{M\pi}-T_{M\lambda})T_{CM}T_{M\lambda}\left[D^{[3]}_{\pi_C}+D^{[3]}_s+T^2_{MC}D^{[3]}_M+D^{[3]}_\lambda\right]+T_{MC}T_{CM}(T_{M\pi}-T_{M\lambda})T_{M\lambda}D^{[1]}_M+T_{CM}T_{M\lambda}D^{[1]}_\lambda$.
4. **Regulation**: $T_RT_{M\lambda}A^0_M+T_{CM}T_RT_{CM}T_{M\lambda}\left[D^{[3]}_{\pi_C}+D^{[3]}_s+T^2_{MC}D^{[3]}_M+D^{[3]}_\lambda\right]+T_{MC}T_{CM}T_RT_{M\lambda}D^{[1]}_M$.

##### 2.3.2 $\chi(\pi_Y, \lambda)$

1. **Catabolism**: $T_{CM}T_{M\lambda}A_s(-t)$.
2. **Dilution**: 0.
3. **Common**: $T_{M\pi}T_{M\lambda}B_M + T_{CM}T_{M\pi}T_{CM}T_{M\lambda}\left[S_{\pi_C} + S_s + T_{MC}^2 S_M + S_\lambda\right]$ $+ 2\frac{\lambda_E}{\lambda_0}T_{M\pi}T_{MC}T_{CM}T_{M\lambda}S_M - T_{CM}T_{M\pi}A_\lambda$.
4. **Regulation**: $T_R T_{M\lambda}B_M + T_{CM}T_R T_{CM}T_{M\lambda}\left[S_{\pi_C} + S_s + T_{MC}^2 S_M + S_\lambda\right]$ $+ 2\frac{\lambda_E}{\lambda_0}T_R T_{MC}T_{CM}T_{M\lambda}S_M - T_{CM}T_R A_\lambda$.

##### 2.3.3 $\chi(\phi_H, \lambda)$

1. **Catabolism**: 0.
2. **Dilution**: $-A_\lambda^0$.
3. **Common**: $(T_{M\pi}-T_{M\lambda})T_{M\lambda}A_M^0+T_{CM}T_{MC}\left(T_{MC}T_{M\lambda}D_M^{[2]} - D_\lambda^{[2]}\right)+T_{CM}(T_{M\pi}-T_{M\lambda})T_{CM}T_{M\lambda}\left[D_{\pi C}^{[3]}+ D_s^{[3]} + T_{MC}^2 D_M^{[3]} + D_\lambda^{[3]}\right] + T_{MC}T_{CM}(T_{M\pi} - T_{M\lambda})T_{M\lambda}D_M^{[1]} + T_{CM}T_{M\lambda}D_\lambda^{[1]}$.
4. **Regulation**: 0.

##### 2.3.4 $\chi(\pi_H, \lambda)$

1. **Catabolism**: 0.
2. **Dilution**: 0.
3. **Common**: $T_{M\pi}T_{M\lambda}B_M+T_{CM}T_{M\pi}T_{CM}T_{M\lambda}\left[S_{\pi_C}+S_s+T_{MC}^2 S_M+S_\lambda\right]+2\frac{\lambda_E}{\lambda_0}T_{M\pi}T_{MC}T_{CM}T_{M\lambda}S_M- T_{CM}T_{M\pi}A_\lambda$.
4. **Regulation**: 0.

#### 2.4 Example: interpretation of the decomposition of the $\phi_Y - \lambda$ cross-covariance

To clarify the interpretation of the expressions above, we now discuss the decomposition of the $Y$–$\lambda$ cross-correlation function in detail.

1. **Catabolism mode**. This mode is a consequence of fluctuations in the expression level of the catabolic sector (C-sector) that directly influence the rate of metabolism and the growth rate. Even though the reporter $Y$ is itself not metabolically active, it does share a noise source ($N_s$) with the C-sector. As a result, the (cross-)correlation $R_{(\phi_Y,\lambda)}$ also contains the catabolism mode, reflecting that $\phi_Y$ correlates with $\phi_C$ (due to the shared noise source $N_s$), and $\phi_C$ with $\lambda$ (due to the effect that fluctuations in $\phi_C$ have on the growth rate).
$$T_{CM}T_{M\lambda}D_s^{[1]}.$$

Note that indeed only the shared noise source $N_s$ transfers directly to growth from $\pi_Y$, instead of reverberating through metabolism (and thus becoming 'common' noise). The function $D_s^{[1]}$ appears (instead of the function $S$ as found in [1]) because noise transferring from metabolism, via production to the reporters ($Y$ or $H$) has a delay timescale of $\lambda_0$, whereas noise transferring to $\phi_C$ has timescale $\lambda_E$.

2. **Dilution mode**. This represents direct transfer from growth to protein concentration.

$$-A_\lambda^0.$$

3. **Common mode**.

$$(T_{M\pi} - T_{M\lambda})T_{M\lambda}A_M^0 + T_{CM}T_{MC}\left(T_{MC}T_{M\lambda}D_M^{[2]} - D_\lambda^{[2]}\right) + T_{CM}(T_{M\pi} - T_{M\lambda})T_{CM}T_{M\lambda}\Big[D_{\pi C}^{[3]}$$
$$+ D_s^{[3]} + T_{MC}^2 D_M^{[3]} + D_\lambda^{[3]}\Big] + T_{MC}T_{CM}(T_{M\pi} - T_{M\lambda})T_{M\lambda}D_M^{[1]} + T_{CM}T_{M\lambda}D_\lambda^{[1]}.$$

Common noise includes noise in the $C$-sector and its effect on the reporters after it reverberated through the cell. Here, one can for example examine the term $(-T_{CM}T_{M\lambda})\,(-T_{CM}(T_{M\pi} - T_{M\lambda}))\,D_\lambda^{[3]}$, where both the path that ends up in $\lambda$, $(-T_{CM}T_{M\lambda})$, and the path that ends up in $\phi_Y$, $(-T_{CM}(T_{M\pi} - T_{M\lambda}))$, cross $\phi_C$, picking up a timescale of $\lambda_E$, and after that one route transfers to $\phi_Y$, picking up a timescale $\lambda_0$. Another interesting example is the second term, $-T_{CM}(T_{M\pi} - T_{M\lambda})D_\lambda^{[2]}$: this is growth noise that diluted $\phi_C$, which has affected metabolism, but from there transfers equally to all other proteins (with coefficient $T_{M\pi} - T_{M\lambda}$). Therefore, this term is also regarded as common noise.

4. **Regulation**

$$T_R T_{M\lambda}A_M^0 + T_{CM}T_R T_{CM}T_{M\lambda}\left[D_{\pi C}^{[3]} + D_s^{[3]} + T_{MC}^2 D_M^{[3]} + D_\lambda^{[3]}\right] + T_{MC}T_{CM}T_R T_{M\lambda}D_M^{[1]}.$$

In the decompositions the term $T_{M\pi} - T_{M\lambda}$ appears often. This represents the net transfer from $M$ to $\phi$ via $\pi$ and from $M$ to $\phi$ via $\lambda$.

#### 2.5 Noise routes, timescales and modes

We make a clear distinction between noise *routes*, *i.e.*, possible paths through which noise can transfer according to the diagram in Fig. S1, and noise *modes*, which are a classification of noise routes in terms of a biophysical noise propagation mechanism (such as dilution or regulatory control). In that sense, a noise mode is a collection of noise routes that each have a similar biological or biophysical interpretation.

The functional forms $A_x^{(0,E)}, S_x, B_x$ and $D_x^{[1,2,3]}$ result from particular noise routes starting at some noise source $N_x$. Here and in Fig. S2, we give some more intuition on how particular noise routes result in the functional forms $D_x^{[1]}$, $D_x^{[2]}$ and $D_x^{[3]}$. First, $D_x^{[1]}$ reflects noise that transfers once via $\phi_C$ and once directly to $\phi_Y$, obtaining timescales $1/\lambda_E$ and $1/\lambda_0$, respectively. An example would be noise source $N_s$ which acts on $\pi_Y$ and therewith directly to $\phi_Y$, but also affects $\phi_C$, transferring to $M$ and further to $\lambda$. As a result $D_x^{[1]}$ is very similar to $S$ [1], which reflect noise routes for which both paths obtain a timescale $1/\lambda_E$. Second, $D_x^{[2]}$ represents a noise source from which one noise route is instantaneous, but the other makes a "double" loop, affecting first $\phi_C$ and picking up timescale $1/\lambda_E$ and then transferring to $\phi_Y$, picking up timescale

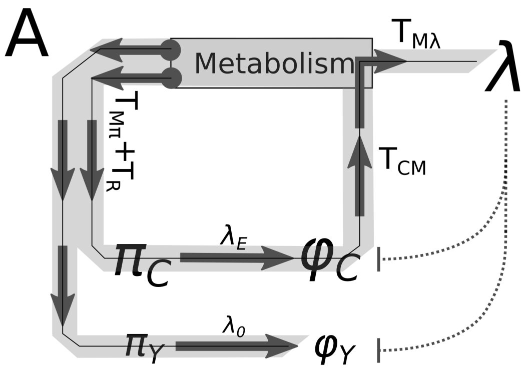

*Figure S2:* Example analysis of two different noise routes from noise source $N_M$ to $\phi_Y$ and $\lambda$ (shown here is a only part of model). Thick gray arrows indicate the direction of a noise route and gray arrows with circles indicate the start of a route. (A) One path goes directly to $\phi_C$ and one path goes, via $\phi_C$ back to $M$ and then transfers to $\lambda$. Transfer coefficients can be split up to represent the paths to $\phi_C$ and to $\lambda$ separately as: $(T_{MC})$ $(T_{MC}T_{CM}T_{M\lambda})$. (B) Another example of a $M$-noise route: $(T_{MC}T_{CM}T_{MC})$ $(T_{M\lambda})$.

$1/\lambda_0$. An example would be noise source $N_M$ which instantaneously affects $\lambda$, but also transfers to $\phi_C$, back to $M$ and then to $\phi_Y$. Third, $D_x^{[3]}$ appears when one route has timescale $1/\lambda_E$ and the other route is "double", picking up both $1/\lambda_E$ and $1/\lambda_0$ timescales. An example of this is another route originating at the noise source $N_s$, where noise first transfers to $\phi_C$, propagates to $M$, and from there to $\lambda$, but also back to $\phi_Y$.

See Fig. S2 for an example, where we compare two routes that originate at noise source $N_M$ in the second line of Eq. 21. A route, mathematically displayed as a product of transfer parameters, is a path through the cell via which noise transfers from the source, in this case $N_M$, to the observables (for example $\phi_Y$ and $\lambda$, as in Fig. S2). One possible route consists of a path directly from $M$, via $\pi_Y$, to $\phi_Y$ (with transfer parameter $(T_{MC})$), and a path from $M$ to $\phi_C$, back to $M$ and onward to $\lambda$ (with transfer parameter $(T_{MC}T_{CM}T_{M\lambda})$, Fig. S2A). This route will contribute a function $D_M^{[1]}$, as one path picks up a timescale of $1/\lambda_0$ and one $1/\lambda_E$

Another possible route consists of a path from $M$ directly to $\lambda$ (transfer parameter $(T_{M\lambda})$), and one route to $\phi_C$, back to $M$, and then to $\phi_Y$ (with transfer parameters $(T_{MC}T_{CM}T_{MC})$). This route will yield a function $D_M^{[2]}$, as one path is instantaneous ($M$ to $\lambda$) and one first picks up a timescale of $1/\lambda_E$ and then one of $1/\lambda_0$.

### 3 Parameter reduction

In this section, we show that two parameters can be effectively scaled away from the system, resulting in fewer parameters to be fitted. First of all, since cross-correlations are dimensionless measures and the equations are linear, we can set one (non-zero) noise amplitude to 1. We pick $\theta_s$, since this noise source has to be present to find any difference between the two reporters in cAMP-fixed cells. Second, note that we do not measure $M$, so that the scale of $M$ is arbitrary and we can scale away $T_{CM}$ and consider fluctuations in $\delta M/(T_{CM}M_0)$ with scaled noise source $N_M' = N_M/T_{\mathrm{CM}}$.

Now, to do the parameter reductions formally, we start with the original system (Eqs. 1-8):

$$\frac{\delta\lambda}{\lambda_0} = N_\lambda + T_{\mathrm{M}\lambda}\frac{\delta M}{M_0}, \tag{39}$$

$$\frac{\delta M}{M_0} = N_{\mathrm{M}} + T_{\mathrm{CM}}\frac{\delta\phi_C}{\phi_{C,0}} = T_{\mathrm{CM}}\left(\frac{N_M}{T_{\mathrm{CM}}} + \frac{\delta\phi_C}{\phi_{C,0}}\right), \tag{40}$$

$$\frac{\delta\pi_C}{\pi_{0,C}} = N_{\pi_{\mathrm{C}}} + N_s + (T_{\mathrm{M}\pi} + T_R)\frac{\delta M}{M_0}, \tag{41}$$

$$\frac{\delta\pi_y}{\pi_{0,y}} = N_{\pi_{\mathrm{Y}}} + N_s + (T_{\mathrm{M}\pi} + T_R)\frac{\delta M}{M_0}, \tag{42}$$

$$\frac{\delta\pi_H}{\pi_{0,H}} = N_{\pi_{\mathrm{H}}} + T_{\mathrm{M}\pi}\frac{\delta M}{M_0}. \tag{43}$$

Defining $\delta\hat{M} = \frac{\delta M}{T_{\mathrm{CM}}M_0}$, $\hat{N}_M = N_M/T_{\mathrm{CM}}$, $\hat{T}_{\mathrm{M}\lambda} = T_{\mathrm{CM}}T_{\mathrm{M}\lambda}$, $\hat{T}_{\mathrm{M}\pi} = T_{\mathrm{CM}}T_{\mathrm{M}\pi}$ and $\hat{T}_R = T_{\mathrm{CM}}T_R$, this system can then be transformed to:

$$\frac{\delta\lambda}{\lambda_0} = N_\lambda + T_{\mathrm{M}\lambda}\left(T_{\mathrm{CM}}\delta\hat{M}\right) = N_\lambda + \hat{T}_{\mathrm{M}\lambda}\delta\hat{M}, \tag{44}$$

$$\delta\hat{M} = \hat{N}_{\mathrm{M}} + \frac{\delta\phi_C}{\phi_{C,0}} \tag{45}$$

$$\frac{\delta\pi_C}{\pi_{0,C}} = N_{\pi_{\mathrm{C}}} + N_s + (T_{\mathrm{M}\pi} + T_R)\left(T_{\mathrm{CM}}\delta\hat{M}\right) = N_{\pi_{\mathrm{C}}} + N_s + (\hat{T}_{\mathrm{M}\pi} + \hat{T}_R)\delta\hat{M}, \tag{46}$$

$$\frac{\delta\pi_y}{\pi_{0,y}} = N_{\pi_{\mathrm{Y}}} + N_s + (T_{\mathrm{M}\pi} + T_R)\left(T_{\mathrm{CM}}\delta\hat{M}\right) = N_{\pi_{\mathrm{Y}}} + N_s + (\hat{T}_{\mathrm{M}\pi} + \hat{T}_R)\delta\hat{M}, \tag{47}$$

$$\frac{\delta\pi_H}{\pi_{0,H}} = N_{\pi_{\mathrm{H}}} + T_{\mathrm{M}\pi}\left(T_{\mathrm{CM}}\delta\hat{M}\right) = N_{\pi_{\mathrm{H}}} + \hat{T}_{\mathrm{M}\pi}\delta\hat{M}. \tag{48}$$

Next, we note that all (cross-co)variances are linear in $\theta_x^2$; hence, if we scale all noise sources by $\theta_s$, the resulting cross-correlations (cross-covariances divided by variances) do not change. How the parameter reduction, and the scaling by $\theta_s$, affects the functional form of the cross-correlation between signals $A$ and $B$ is shown below, where we split the cross-covariance and the variances in contributions of each noise source. Let the entire contribution (written in hat-parameters) of each noise source $x$ to the cross-covariance between signals $A$ and $B$ be written as $\theta_x^2 f_x$ (time dependence is omitted for readability), and write $\theta_x^2 g_{A,x}$, and $\theta_x^2 h_{B,x}$, for the contribution of noise source $N_x$ to the variance of $A$ and $B$ respectively. Then:

$$R_{(A,B)}(\tau) = \frac{\left(\frac{\theta_M}{T_{\mathrm{CM}}}\right)^2 f_{\hat{M}} + \sum_{i\neq M}\theta_i^2 f_i}{\sqrt{\left(\frac{\theta_M}{T_{\mathrm{CM}}}\right)^2 g_{A,\hat{M}} + \sum_{i\neq M}\theta_i^2 g_{A,i}}\sqrt{\left(\frac{\theta_M}{T_{\mathrm{CM}}}\right)^2 h_{B,\hat{M}} + \sum_{i\neq M}\theta_i^2 h_{B,i}}} \tag{49}$$

$$= \frac{\left(\frac{\theta_M/\theta_s}{T_{\mathrm{CM}}}\right)^2 f_{\hat{M}} + f_s + \sum_{i\neq s,M}(\theta_i/\theta_s)^2 f_i}{\sqrt{\left(\frac{\theta_M/\theta_s}{T_{\mathrm{CM}}}\right)^2 g_{A,\hat{M}} + g_{A,s} + \sum_{i\neq s,M}(\theta_i/\theta_s)^2 g_{A,i}}\sqrt{\left(\frac{\theta_M/\theta_s}{T_{\mathrm{CM}}}\right)^2 h_{B,\hat{M}} + h_{B,s} + \sum_{i\neq s,M}(\theta_i/\theta_s)^2 h_{B,i}}} \tag{50}$$

$$= \frac{\left(\hat{\hat{\theta}}_M\right)^2 f_{\hat{M}} + f_s + \sum_{i\neq s,M}\hat{\theta}_i^2 f_i}{\sqrt{\left(\hat{\hat{\theta}}_M\right)^2 g_{A,\hat{M}} + g_{A,s} + \sum_{i\neq s,M}\hat{\theta}_i^2 g_{A,i}}\sqrt{\left(\hat{\hat{\theta}}_M\right)^2 h_{B,\hat{M}} + h_{B,s} + \sum_{i\neq s,M}\hat{\theta}_i^2 h_{B,i}}}. \tag{51}$$

Here, $\hat{\theta}_x = \theta_x/\theta_s$, and $\hat{\hat{\theta}}_M = \frac{\theta_M/\theta_s}{T_{\rm CM}}$. By considering only $\hat{\hat{.}}$ - parameters, we effectively removed two parameters from the model.

### 4 Data analysis

#### 4.1 Calculating cross-correlations from data

Segmentation was done using the software Schnitzcells, developed by the Elowitz lab [3], with custom scripts written by Daan Kiviet, Philippe Nghe and Noreen Walker [1, 4]. Tracking was done in line with Kiviet *et al* [1]. Cell length is determined by fitting a 3rd (or, in some cases 5th) order polynomial through the cell area. Cross-covariance $\chi$ and cross-correlation $R$ between two signals in discrete time is then defined as:

$$\chi_{f,g}(\tau) = \frac{1}{N-1} \sum_{n=0}^{N-|\tau|-1} \hat{f}(n)\hat{g}(n+\tau). \tag{52}$$

$$R_{f,g}(\tau) = \frac{S_{f,g}(\tau)}{\sqrt{S_{f,f}(0)S_{g,g}(0)}}. \tag{53}$$

Here, hats indicate mean-subtracted signals.

Cells that are born earlier in the experiments appear in more lineages. When calculating cross-correlations along lineages, we must thus be careful to not count such cells repeatedly. Therefore, we introduce for each data point a weight, representing in how many branches the point occurs. The resulting cross-correlation is a composite cross-correlation with contributions of points from multiple branches $i$. Lastly, we also introduce time-average-subtracted variable, since averages can change slightly during experiments. We therefore define a composite cross-covariance and cross-correlation:

$$\mathcal{S}_{f,g}(\tau) := \frac{1}{W_{\text{total},\tau}} \sum_i \frac{1}{N_i - |\tau|} \sum_{n=0}^{N_i-|\tau|} w_{n,i,\tau}\hat{f}_i(n)\hat{g}_i(n+\tau), \tag{54}$$

$$\mathcal{R}_{f,g}(\tau) = \frac{\mathcal{S}_{f,g}(\tau)}{\sqrt{\mathcal{S}_{f,f}(0)\mathcal{S}_{g,g}(0)}}, \tag{55}$$

$$\hat{f}_i(n) = f_i(n) - \langle f_i \rangle_n \tag{56}$$

$$w_{n,i,\tau} = 1/K_{n,i,\tau}, \tag{57}$$

$$W_{\text{total},\tau} = \sum_{n,i} w_{n,i,\tau}. \tag{58}$$

Here, the summations run are over all branches $i$ and time points $n$. Weights are indicated with $w$, where $K_{n,i,\tau}$ is the frequency with which a specific point pair $\hat{f}_i(n)\hat{g}_i(n+\tau)$ was used. Throughout the manuscript we refer to the composite cross-correlation $\mathcal{R}$ as the cross-correlation $R$. The mean-subtracted signal $\hat{f}_i(n)$ is now recalculated in each branch, for each time point to compensate for a changing overall average during the experiment.

#### 4.2 Averaging multiple experiments and estimating error bars

To create Figs. 2 and 3 of the main text, the cross-correlations calculated per microcolony were averaged. Here, we explain how we averaged the multiple (independent) experiments (each microcolony being an experiment) and how we calculated error bars.

We use a Mixed Model Estimate, assuming that each individual measurement $y_{i,j}$ from experiment $i$ is determined by the average of interest, $\mu$, plus two noise sources: within experimental noise $\xi_j(i)$, and between-experiments noise $\xi_i$:

$$y_{i,j} = \mu + \xi_i + \xi_j(i).$$

Here, $\xi_i$ is the noise that determines the off-set of the mean from experiment $i$, and $\xi_j(i)$ the noise on individual measurement $j$ in group $i$. We assume that $\mathbb{E}[\xi_i] = \mathbb{E}\left[\xi_j\right] = 0$. Furthermore, let $\text{Var}\left(\xi_j\right) = s_i^2$, the variance within experiment $i$ (this variance might differ between experiments), and $\text{Var}(\xi_i) = s_\mu^2$, the variance between the means of each experiment. The within experiment variance is estimated by dividing each microcolony into four lineages (from the moment there were four cells in the microcolony, we followed each of their lineages separately) and calculating and comparing statistics along each lineage. With these notions for $\xi_i$ and $\xi_j(i)$, we write $n$ for the number of experiments and $n_i = 4$ for the number of subgroups within experiment $i$. Then:

$$\mathbb{E}\left[y_{i,j}\right] = \mu \tag{59}$$

$$\text{Var}\left(\mathbb{E}\left[y_{i,j}\right]\right) = \frac{\text{Var}\left(\sum_i \sum_j \mu + \xi_i + \xi_j(i)\right)}{\left(\sum_i n_i)^2\right)} \tag{60}$$

$$= \frac{1}{\left(\sum_i n_i\right)^2}\text{Var}\left(\sum_i n_i \xi_i + \sum_i \sum_j \xi_j\right) \tag{61}$$

$$= \frac{1}{\left(\sum_i n_i\right)^2}\left(\sum_i n_i^2 \text{Var}\left(\xi_i\right) + \sum_i n_i \text{Var}\left(\xi_j\right)\right) \tag{62}$$

$$= \frac{s_\mu^2 \sum_i n_i^2 + \sum_i n_i s_i^2}{\left(\sum_i n_i\right)^2} \tag{63}$$

To estimate $\mu$, we again use knowledge of within-experiment variances to calculate a weight factor for each microcolony: let $y_i(t)$ be the measured mean value of an observable in experiment $i$ at time $t$, with within-experiments error $s_i(t)$. Then we estimate $\mu$ as:

$$\langle y(t)\rangle = \frac{\sum_i^n s_i^{-2}(t) y_i(t)}{\sum_j^n s_j^{-2}(t)} =: \frac{\sum_i w_i(t) y_i(t)}{W(t)}.$$

Here, $w_i(t) = \frac{1}{s_i^2(t)}$ and $W(t) = \sum_{j=1}^n w_i(t)$. That is, more precise measurements (that is, those with smaller within-experiment error $s_i$) obtain a higher weight.

#### 4.3 Null-expectation for the cross-correlations

To confirm that the measured cross-correlations correspond to real, biological, signals inside the cells, we performed a permutation analysis on the time-series data. We kept the temporal information of the data, but randomized at each time point the growth rate and expression data for all the cells in the colony. Any biological correlations between variables should therewith be removed. Repeating this randomization 50 times, and each time re-calculating cross-correlations, indeed gives a band of cross-correlations around zero, allowing us to infer what kind of signals could still be explained purely by technical noise (See for example Fig. S9). Any part of the originally measured cross-correlations that fall outside this band can then be concluded to stem from a real biological signal.

### 5 Fitting procedure

#### 5.1 Parameter values for *WT* and *cAMP-fixed** cells

The full mathematical model, with the reduced number of parameters, was fitted to the cross-correlations with their error bars using a weighted least-square fitting procure in Mathematica 13. We fitted $R_{(\phi,\lambda)}$ and $R_{(\pi,\lambda)}$ for both reporters and for wild type and cAMP-fixed* cells simultaneously.

In all fits, we set $\theta_{\pi_C} = 0$, for three reasons. First, the variable $\phi_C$ represents the entire C-sector, so that random, intrinsic fluctuations in the total size of this sector are expected to be small. Second, any noise source that directly influences the total size of the C-sector should also affect the CRP-reporter Y, because C and Y are regulated and expressed similarly. Such noise sources are therefore also captured by the phenomenological noise source $N_s$ that summarizes shared noise sources between the C-sector and Y. Third, fluctuations in the concentrations of each of the individual proteins of the C-sector likely transfer differently to metabolism than joined fluctuations of the entire C-sector. The effect of the intrinsic fluctuations of individual C-sector protein species on the metabolic flux is instead captured by the noise source $N_M$. Fluctuation in any individual C-sector protein could indeed potentially influence the flux catalyzed by the entire C-sector, causing fluctuations in metabolism.

For cAMP-fixed* cells we predefined that $T_R = 0$, and for wild type cells $T_R < 0$. All other parameters were not allowed to differ between WT and cAMP-fixed* cells. We thus fitted 9 free parameters: $\{\hat{T}_R$ (only for WT cells), $\hat{T}_{\mathrm{M}\lambda}, \hat{T}_{\mathrm{M}\pi}, \hat{\theta}_{\pi_Y}, \hat{\theta}_{\pi_h}, \hat{\hat{\theta}}_M, \hat{\theta}_\lambda, \frac{\beta_\pi}{\lambda_0}, \frac{\beta_\mu}{\lambda_0}\}$. (The hat-parameters are defined in the section‘Parameter Reduction’, section 3; $T_{\mathrm{CM}}$ and $\theta_s$, are set to unity and removed from the model by scaling.) The timescale $1/\beta_\pi$ is the time-scale of the production noise rates, and $1/\beta_\mu$ is the timescale of growth/dilution-related noise sources ($\theta_s$, $\theta_M$, $\theta_\lambda$). Their value given is relative to the mean growth rate, $\lambda_0$.

In table S1 we present best-fit parameters, with their 95% confidence interval. The 95% confidence ranges of the parameters were estimated by changing that parameter until the increase in (weighted) sum of squared residuals was statistically significant (as determined using $F$-statistics).

Although the model is able to reproduce the experimentally observed cross-correlations, we are hesitant to over-interpret the exact numerical values of the fitted parameters. For example, the wild type and optimal mutant's best-fit-parameter for the transfer from $M$ to $\pi$, $\hat{T}_{\mathrm{M}\pi}$, is small, but negative (Table S1). Such a negative parameter is counter-intuitive, for it results in a negative common mode (upward fluctuations in metabolism increase the growth rate, but lower the production rates). Presence of this negative common mode is only reflected in the mutant's negative $\pi_H$-$\lambda$ cross-correlation, and, importantly, it is also this negative cross-correlation that causes the best-fit parameter to become negative (not including $R_{(\pi_H,\lambda)}$ causes the best-fit value for $\hat{T}_{\mathrm{M}\pi}$ to be slightly positive, data not shown). Extensions of the model with alternative mechanistic explanations that could explain the negative correlation between $\pi_H$ and $\lambda$ (such as competition between mRNA molecules for ribosomal binding sites at the single-cell level, or a negative effect of the reporters on the growth rate [5]) did also not fit the negative cross-correlation well (data not shown, Mathematica notebooks available upon request with the authors). Possibly, this specific negative cross-correlation could be caused by an experimental artifact that heavily influences the fitted parameters. Indeed, similar constitutive reporters measured in earlier work [1, 6] show a positive correlation between $\pi_H$ and $\lambda$.

| Parameter | Confined to | Best Fit | 95% Confidence Interval |
|---|---|---|---|
| $\theta_{\pi_C}$ | 0 | - | - |
| $\theta_s$ | 1 | - | - |
| $T_{\mathrm{CM}}$ | 1 | - | - |
| $T_R$ | $[-2, 2]$ ($WT$); 0 ($MUT$) | **-1.67 (WT)** | [-2.28,-1.25] |
| $\hat{T}_{\mathrm{M}\lambda}$ | $[-1, 1]$ | **0.05** | [0.045, 0.54] |
| $\hat{T}_{\mathrm{M}\pi}$ | $[-1, 1]$ | **-0.179** | [-0.22, -0.14] |
| $\hat{\theta}_{\pi_Y}$ | $[0, 5]$ | **0.0** | [0,0.74] |
| $\hat{\theta}_{\pi_h}$ | $[0, 5]$ | **1.14** | [1.06,1.24] |
| $\hat{\hat{\theta}}_M$ | $[0, 10]$ | **0.90** | [0.83,0.97] |
| $\hat{\theta}_\lambda$ | $[0, 5]$ | **0.13** | [0.125,0.143] |
| $\beta_\pi$ | $> 2$ | **2** | [1.8, 2.16] |
| $\beta_\mu$ | $> 0.5$ | **0.78** | [0.75,0.82] |

*Table S1:* Best fit parameters of the wild type and optimal mutant, including predefined parameter constraints and best-fit confidence ranges based on statistical analysis. Parameter $T_R$ was set to 0 for *cAMP-fixed** ($MUT$) cells. All other parameters were fitted using a minimization of the squared distance of 8 analytical curves (cross-correlations $\phi - \lambda$ and $\pi - \lambda$ for both reporters. and for $WT$ and *cAMP-fixed** cells) to the cross-correlation data (Fig. 2B from the main text). For the precise interpretation of $\hat{\cdot}$-parameters, see section 'Parameter Reduction'.

#### 5.2 Low and High cAMP

From the best-fit parameters for the cAMP-fixed* cells (Table S1), we qualitatively reproduced the Low cAMP cross-correlations by increasing transfer to, and from, metabolism. Intuitively, when cAMP concentration is low and C-sector expression is sub-optimal, one would expect strong transfer from $\phi_C$ to $M$ (*i.e.* an increase in the parameter $T_{CM}$). Additionally, $M$ -itself being limited by $\phi_C$- is in turn expected to be limiting for growth and protein production; as soon as metabolism becomes better, the cell can grow and create protein faster (corresponding to an increase in parameters $T_{M\pi}$ and $T_{M\lambda}$). Therefore, we first scaled $T_{CM}$ by a factor of 1.1. Note that scaling $T_{CM}$ also affects some of the scaled (hat) parameters; $\hat{T}_{\mathrm{M}\lambda}$, $\hat{T}_{\mathrm{M}\pi}$, and $\hat{\hat{\theta}}_M$. We assumed $\hat{\hat{\theta}}_M$ to follow the scaling (*i.e.* the $\hat{\hat{\theta}}_M$ becomes a factor of 1.1 smaller). The parameters $\hat{T}_{\mathrm{M}\lambda}$ and $\hat{T}_{\mathrm{M}\pi}$, whose values are expected to increase also independently of $T_{CM}$, were instead both increased with a constant, 0.3. This resulted in, for Low cAMP-fixed cells in: $\{\hat{T}_{\mathrm{M}\lambda} = 0.35, \hat{T}_{\mathrm{M}\pi} = 0.121, \hat{\hat{\theta}}_M = 0.82\}$. The timescales of each noise source relative to the growth rate (which is lower under this condition than under optimal cAMP levels) were kept constant.

The resulting cross-correlations reproduce many qualitative features of the measurements. That said, the mathematical model slightly over-estimates the overall amplitude of the cross-correlation (Fig. S6A). Possibly, this is due to changing average expression, or possibly caused by experimental error: Independent measurement noise in any two variables reduces their correlation. In the model, this can be mimicked by increasing the noise levels of $\theta_{\pi_Y}$ and $\theta_{\pi_H}$ to $\{5, 2.14\}$ respectively. An increase in those noise parameters does not affect the shape of the cross-correlation, but only decreases the overall amplitude of the cross-correlation (decorrelation).

For the *cAMP-fixed*$^{\mathrm{high}}$ cells, we assumed that $\phi_C$ now negatively influences (hinders) metabolism, but that a better or faster metabolism still results in faster growth and protein expression. Therefore, we scaled $T_{CM}$ by $-0.3$, in line with its hypothesized slightly negative burden to metabolism and growth. Parameters $\hat{T}_{\mathrm{M}\lambda}$ and $\hat{T}_{\mathrm{M}\pi}$ therewith picked up a minus sign, but we kept their amplitudes as for the *cAMP-fixed*$^{\mathrm{low}}$ cells, since metabolism is still far from optimal, so that noise from metabolism is expected to strongly transfer to growth and protein production. Thus, parameters for the $\mathrm{cAMP-fixed}^{\mathrm{high}}$ cells were: $\{\hat{T}_{\mathrm{M}\lambda} = -0.35, \hat{T}_{\mathrm{M}\pi} = -0.121, |\hat{\hat{\theta}}_M| = 3\}$.

These parameters indeed qualitatively explained many featured of the measured cross-correlations (Fig. S6B). However, the high-cAMP cross-correlations are, as in the low cAMP condition, overestimated by the model. Another clear mismatch is that, for these parameters, the model predicts a strong common mode (which can be seen most clearly in the $\pi_H$-$\lambda$ cross-correlation) that does not seem to be present in the data (although one microcolony did show a clear positive $\pi_H$-$\lambda$ cross-correlation, see bottom right panel in Fig. S7). Interestingly, just as in the wild type, the cross-correlations of both reporters look similar in cAMP-fixed cells under the high-cAMP condition.

A semi-quantitative fit can made (see also Fig. 3E of the main text) by further tuning the reporter's noise amplitudes to: $\{\theta_{\pi_Y} = 2, \theta_{\pi_H} = 3.144\}$.

### 6 Toy model of the means of the two reporters

From the observation that the sum of the concentration of both reporters is, on average (Fig. S10A), approximately constant, we were inspired to also write equations for the population-level average behavior. We here derive a phenomenological toy model that describes how the average concentrations, production rates and growth rates of cAMP-fixed cells change under changing external cAMP concentration.

Generally, we can write for the (average) total concentration of the catabolic sector $\phi_C$ and for the constitutive reporter $H$:

$$\frac{\partial \phi_C}{\partial t} = \pi_C - \phi_C \lambda = (f_C(x) - \phi_C)\lambda, \tag{64}$$

$$\frac{\partial \phi_H}{\partial t} = \pi_H - \phi_H \lambda = (f_H - \phi_H)\lambda. \tag{65}$$

Here $f_C(x)$ is a regulation function that determines the fraction of all metabolic flux allocated towards production of $\phi_C$ as a function of some internal metabolite (in this case, $x$ reflect the internal cAMP concentration). However, $f_H$ is also not necessarily a constant and can depend on resource allocation. We will show below that $f_H$ is indeed not constant.

From the experiments we observe $\phi_Y + \phi_H$ is approximately a constant. Assuming that $Y$ is a good reporter of the $C$ sector, this is equivalent to:

$$\phi_C + \phi_H = T$$

(Note that this ignores proteomic shifts that result from a changing the ribosomal sector, or any other sector that is not modeled here.) In the steady state, $\phi_H = f_H$, and therefore also $\pi_H = \phi_H \lambda = (T - f_C(x))\lambda = f_H \lambda$. We can thus rewrite the differential equation for $H$ as:

$$\frac{\partial \phi_H}{\partial t} = (T - f_C(x))\lambda - \phi_H \lambda.$$

In steady state, this suggest that the production rates (as directly quantified from the experiments), divided by the growth rates should be equal to the concentration for both the reporters (Fig. S10A).

In the mutant, the growth rate declines when either over, or under-expressing $\phi_C$, and the mean growth rates fit a 2nd order polynomial nicely.

$$\lambda(\phi_C) := g_\lambda(\phi_Y) = a\phi_Y^2 + b\phi_Y + c$$

For best-fit parameters $\{a = -3.94 \cdot 10^{-5}, b = 2.12 \cdot 10^{-2}, c = -2.02\}$. Here we consider $\phi_C$, the metabolic sector, well-reported by the metabolic reporter $\phi_Y$.

Using this polynomial, the relationships between mean protein production rates/mean protein concentrations, and the mean growth rate can be related (Figs. S3 and S10):

$$\{\phi_Y, \lambda\} = \{\phi_Y, g_\lambda(\phi_Y)\}, \tag{66}$$
$$\{\pi_Y, \lambda\} = \{\phi_Y g_\lambda(\phi_Y), g_\lambda(\phi_Y)\}, \tag{67}$$
$$\{\phi_H, \lambda\} = \{\phi_H, g_\lambda(T - \phi_H)\}, \tag{68}$$
$$\{\pi_H, \lambda\} = \{\phi_H g_\lambda(T - \phi_H), g_\lambda(1 - \phi_H)\}, \tag{69}$$
$$\{\pi_Y, \pi_H\} = \{\phi_Y g_\lambda(T - \phi_H), \phi_H g_\lambda(T - \phi_H)\}. \tag{70}$$

These relationships fit strikingly well (Fig. S10).

### 7 Supplementary Figures

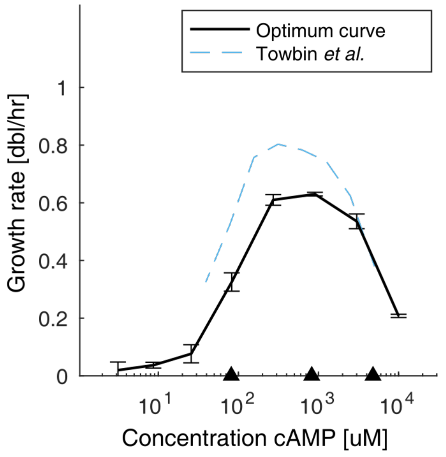

*Figure S3:* Growth rates in minimal medium supplemented with lactose and various cAMP concentrations. Measured exponential phase growth rates of the *cyaA cpdA* null mutant (cAMP-fixed cells) at different concentrations of cAMP as measured in a plate reader. Black triangles refer to the low, optimal and high cAMP concentrations respectively. The growth rate was determined as an exponential fit over a manually selected part of the bacterial density curve. Additionally, this figure shows data from a similar experiment performed by Towbin *et al.* [7]

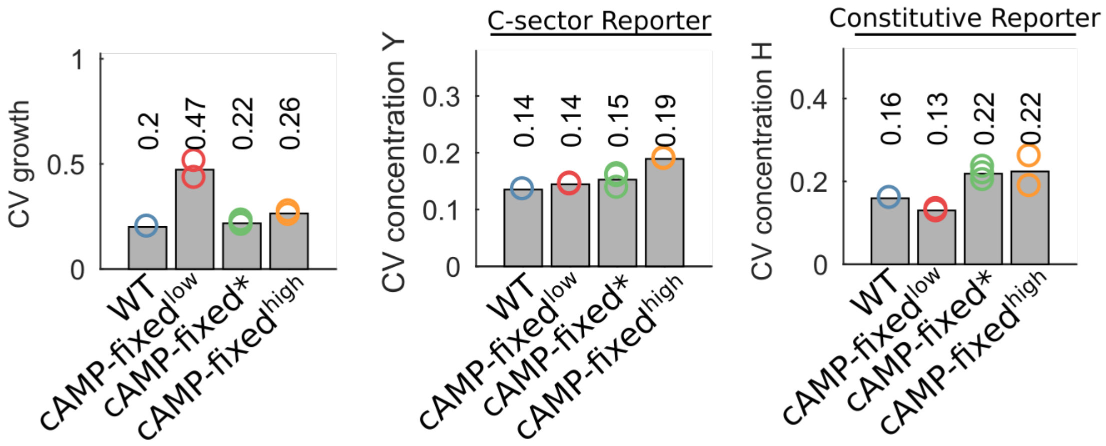

*Figure S4:* Coefficients of variation of the growth rate (left panel) and the concentrations of the C-sector reporter (middle panel) and the constitutive reporter (right panel) for the different conditions. Shown in the figure are only the experiments performed with a similar microscope such that their absolute values were comparable.

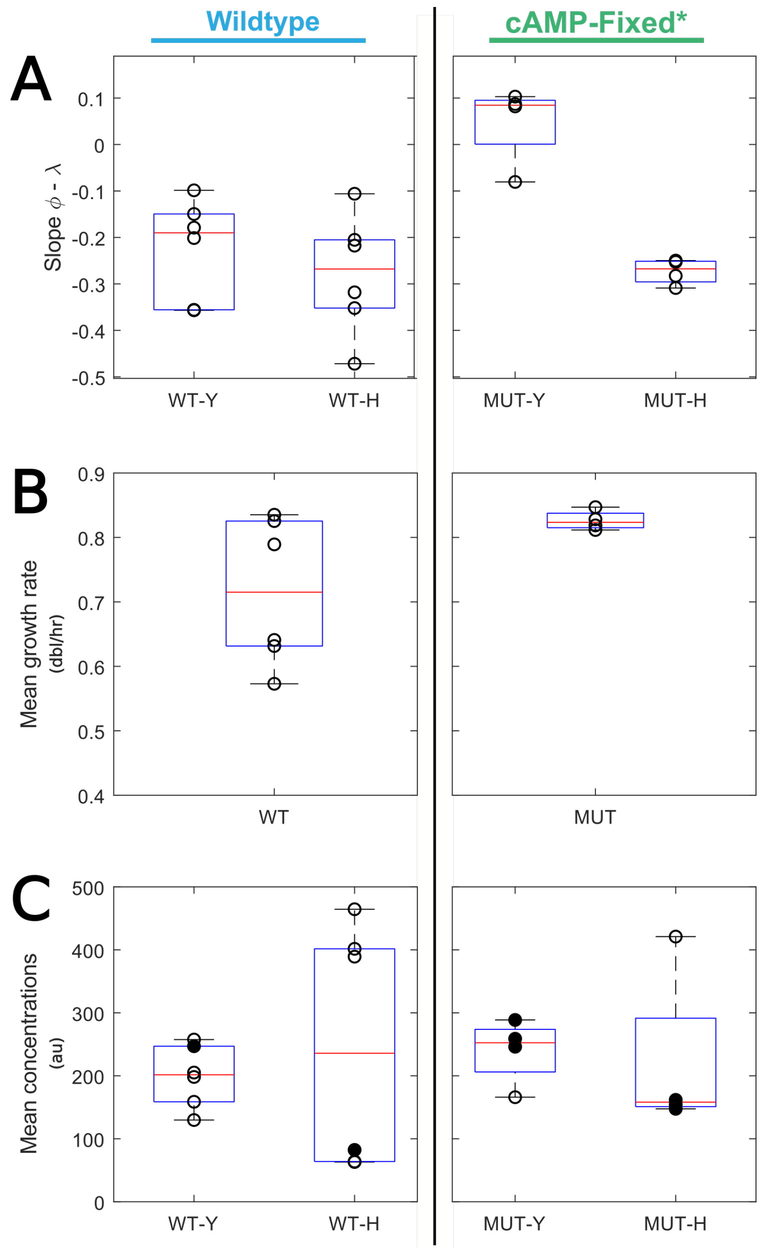

*Figure S5:* Average experimental values measured in different colonies. (**A**) Regression slope between $\phi$ and $\lambda$ for the wild type and *cAMP-fixed** cells (MUT in the figure). Slopes for the *Y* reporter differ significantly between wild type and mutant cAMP-fixed* cells ($p = 0.0031$, two-sample $t$-test), but not for the H reporter ($p = 0.93$, Welch's $t$-test). Note that to calculate these regression slopes only relative fluctuations are relevant, so that only relative fluorescence signals are relevant. (**B**) Average growth rate per colony, showing a large variance in growth-rate measurements. Difference between the mean growth rates of *WT* and *cAMP-fixed** cells is not significant ($p = 0.063$, Welch's $t$-test). (**C**) Average fluorescence per colony. Filled circles are values measured with a standardized microscope setting (and thus only those absolute values can be compared). Y: C-sector reporter, H: constitutive reporter.

| Figures with data | Condition | # colonies | # cells | # data points |
|---|---|---|---|---|
| 1C, 1E | WT | 1 | 1671 | 5979 |
| 1D, 1F | Fixed-800 | 1 | 1580 | 4339 |
| 2B (left panels) | WT | 6 | 113,63,110,953,729,1667 | 573,346,513,2420,1834,5979 |
| 2B (right panels) | Fixed-800 | 4 | 1568, 1626,1500,837 | 4339, 5202,4697,2664 |
| 3A,B,D | Fixed-80 | 2 | 1165, 623 | 4141,1985 |
| 3A,B | Fixed-800† | 3 | 1568, 1626,1500 | 4339, 5202,4697 |
| 3A,B,E | Fixed-5000 | 2 | 1437,837 | 4229, 2086 |

*Table S2:* Number of cells (as determined by the Schnitzcells software (see Methods), number of colonies, and number of data points from all single cell experiments. (†Only a subset of data was used, as for those data sets microscopy conditions were equal, such that the values can be compared directly.)

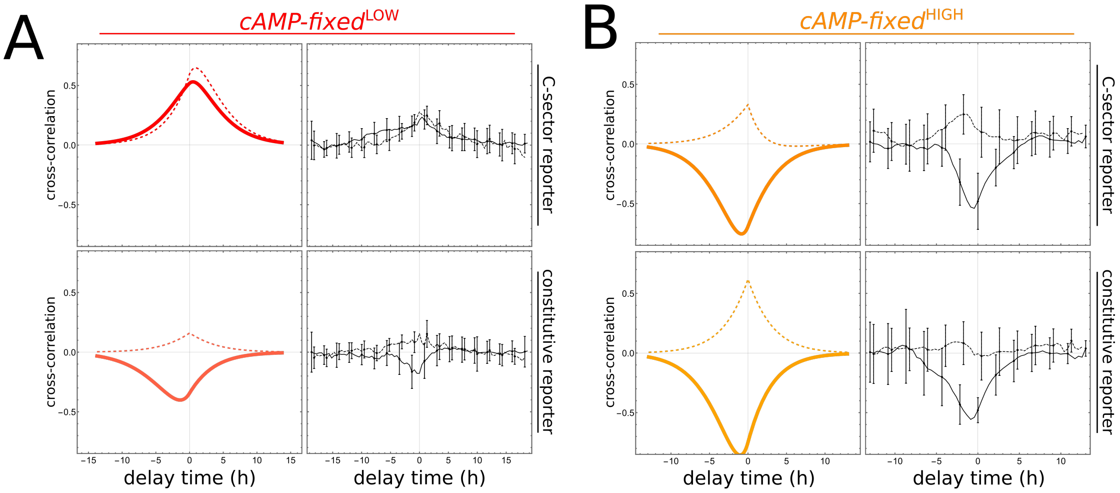

*Figure S6:* Qualitative model prediction for the cross-correlations as measured in the cAMP-fixed cells using low-cAMP (80 μM, panel A) and high-cAMP (5000 μM, panel B) conditions, together with the measured cross-correlations. (A) The transfer parameters from $M$ to $\lambda$ ($\hat{T}_{M\lambda}$) and from $M$ to $\pi$ ($\hat{T}_{M\pi}$) have been increased by 0.3, and $T_{CM}$ is slightly increased as well (scaled by 1.1; affecting affects $\hat{\theta}_M$), compared to the best-fit values from the *cAMP-fixed** condition. (B) Same values as in (A) for $\hat{T}_{M\lambda}$ and $\hat{T}_{M\pi}$, but now $T_{CM}$ is slightly negative (multiplied by −0.3).

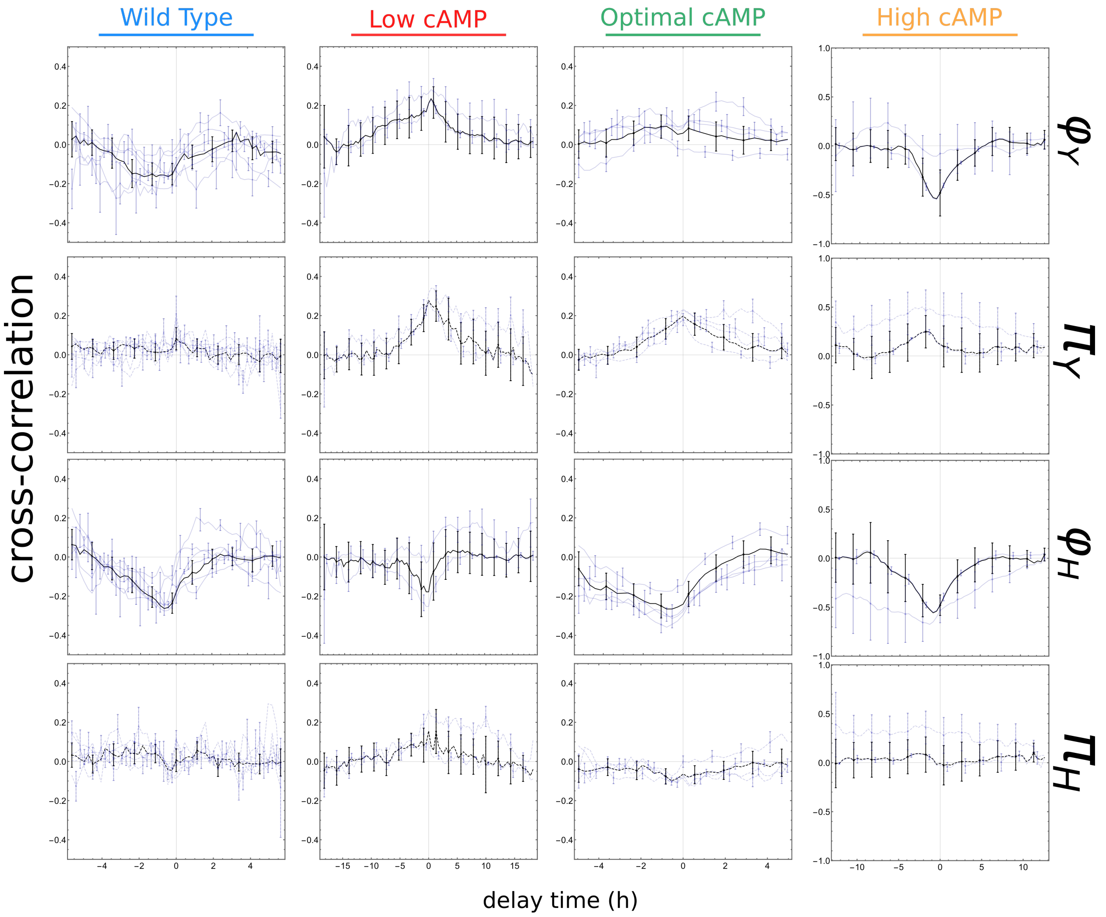

*Figure S7:* All measured cross-correlations (thin blue lines) in independent replicates (independent colonies), together with their weighted averages (thick black lines), for all conditions. Y: C-sector reporter, H: constitutive reporter.

| **Identifier** | **Manuscript shorthand** | **Description** |
|---|---|---|
| ASC838 | | Wild type MG1655, also known as strain bBT12 and CGSC number 8003. Known mutations: $\lambda-$, Δfnr-267, rph-1. (No resistance modules.) |
| ASC839 | | *cyaA*, *cpda* null mutant. Also known as strain bBT80. Based on ASC838. (No resistance modules.) |
| ASC990 | wild type (WT) | Wild type strain, except for Δ(galk)::nCRPr-mCerulean-kanR and Δ(intc)::CRPr-mVenus-cmR. (Kanamycin and chloramphenicol resistant.) |
| ASC1004 | cAMP-fixed | Strain based on ASC839 (ΔcyaA Δcpda), introduced Δ(galk)::s70-mCerulean-kanR and Δ(intc)::rcrp-Venus-cmR. (Kanamycin and chloramphenicol resistant.) |

*Table S3:* Additional details on the strains that were used; ASC990 and ASC1004 were used in the manuscript, and are based on ASC838 and ASC839 respectively.

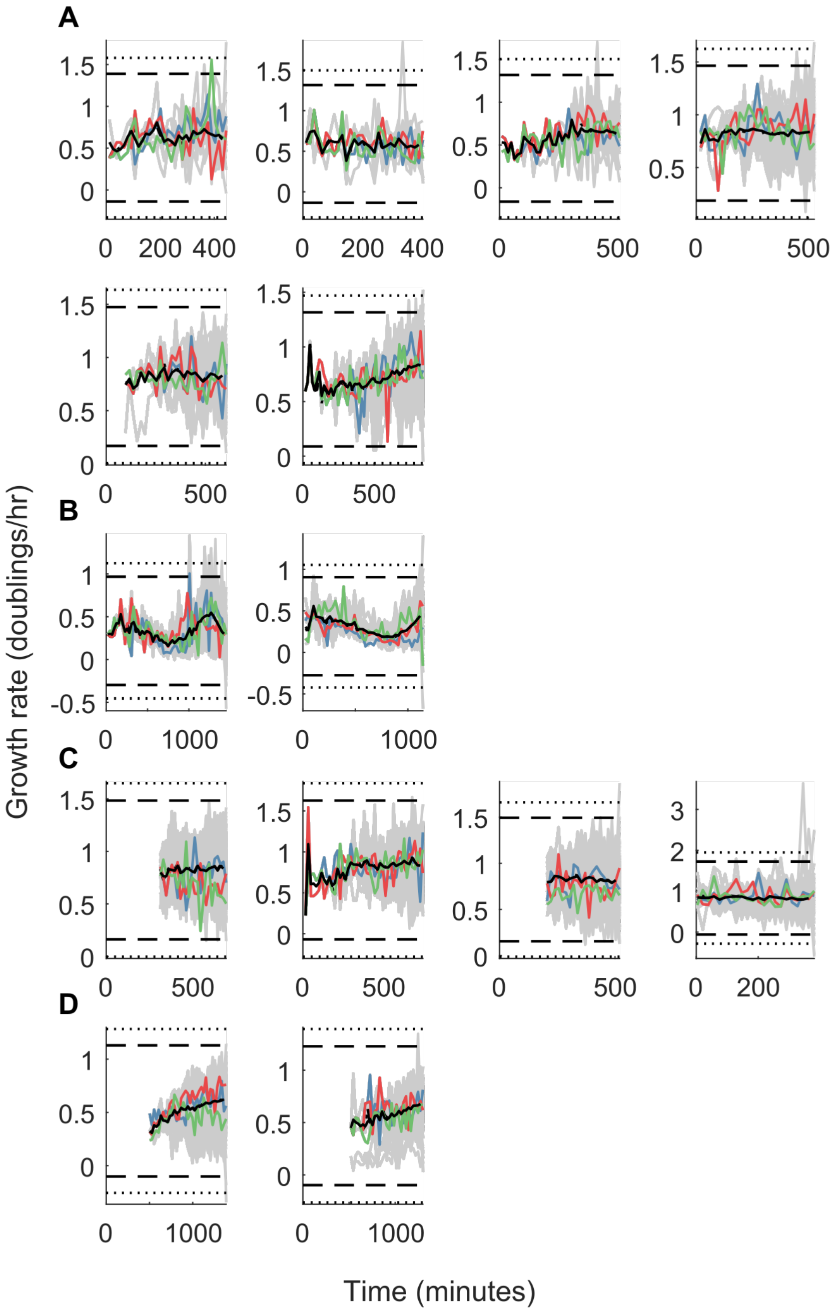

*Figure S8:* Growth rates during the experiments. Each panel plots growth-rate data for a single colony; panels are grouped by growth condition. The gray lines show single lineage traces, the black lines the population average. Colored lines highlight example single lineage traces to illustrate single cell behavior. Dashed and dotted lines indicate $4\sigma$ and $5\sigma$ boundaries from the overall mean respectively. As before, the displayed conditions are (A) wild type cells, (B) cAMP-fixed* cells growing on 80 μM cAMP, (C) cAMP-fixed cells growing on 800 μM cAMP and (D) mutant cells growing on 5000 μMcAMP.

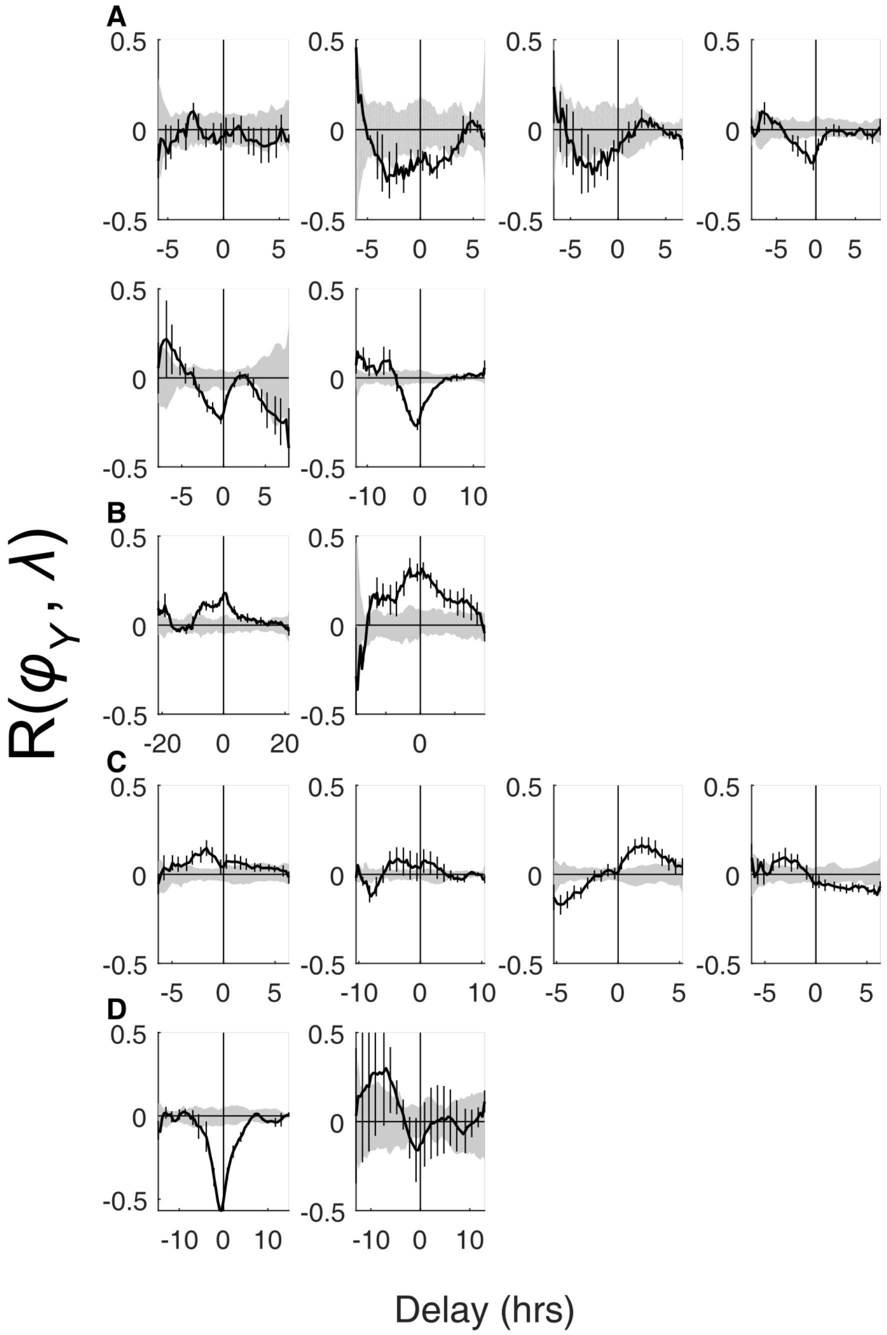

*Figure S9:* Cross-correlations between the C-sector reporter, $\phi_Y$, and $\lambda$, together with their null expectation (gray areas around 0, see section 4.3 for details of the calculation). The black lines in this figures are the light-blue lines in top panels of Figure S7. Error bars are calculated by dividing each microcolony into four parts and comparing statistics in each part. As before, the displayed plots are from independent microcolonies growing under the following conditions: (A) wild type cells, (B) cAMP-fixed cells growing on 80 µM cAMP, (C) cAMP-fixed cells growing on 800 µM cAMP and (D) cAMP-fixed cells growing on 5000 µM cAMP.

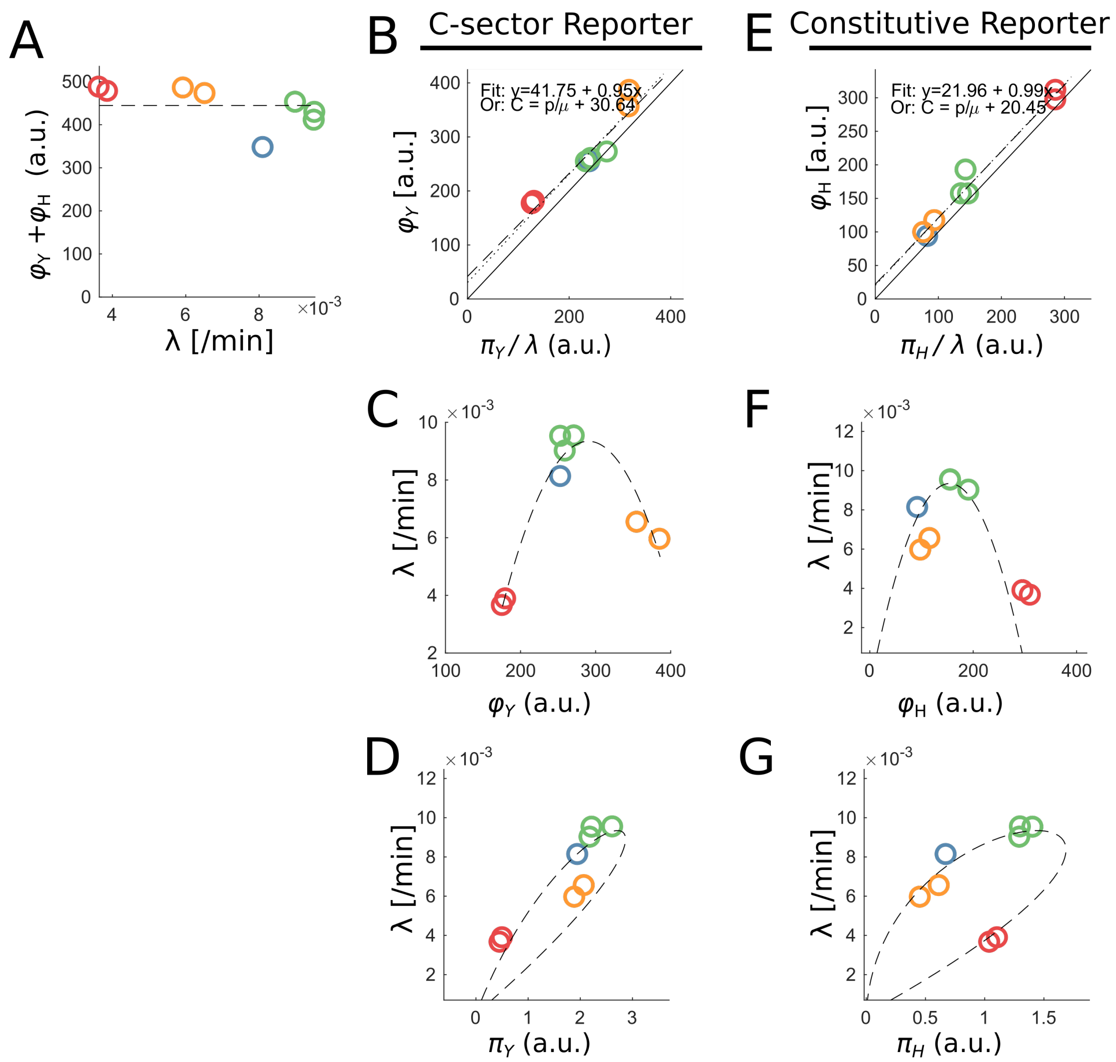

*Figure S10:* Toy model fits the mean behavior of the reporters and the growth rate (see for details section 6). (A) The sum of reporter concentrations is approximately constant in all conditions. (B) Steady state relationship $\phi_Y = \pi_Y/\lambda$ (black straight line) holds closely for all conditions. The best fit (dashed line), however, has a slight offset. (C) Fitted parabolic relationship of $\phi_C$ between the growth conditions. See also S3). (D) Relationship between $\pi_Y$ and $\lambda$ as calculated from the toy model (6). (E) Steady state relationship for the C-sector reporter. (F-G) The C-sector reporter concentrations and production rates for each condition fall on the curve calculated from the toy model (not fitted). Colour coding is as in other figures (blue: wild type, red: low cAMP mutant, green: medium cAMP mutant, orange: high cAMP mutant). Y: C-sector reporter, H: constitutive reporter.

```
PLAC.   GGTTTCCCGACTGGAAAGCGGGCAGTGAGCGCAACGCAATTAATGTGAGTTAGCTCACTC
CRPr.   -----CGTCAGGAGGAGAGGGGCAGTGAGCGCAACGCAATTAATGTGAGTTAGCTCACTC
nCRPr.  -----CGTCAGGAGGAGAGGGGCAGTGAGCGCAACGCAATCAGATCAAATGTGTCGTTTC

pLAC.   ATTAGGCACCCCAGGCTTTACACTTTATGCTTCCGGCTCGTATGTTGTGTGGAATTGTGA
CRPr.   ATTAGGCACCCCAGGCTTTACACTTTATGCTTCCGGCTCGTATGTTGTGTGCATGGATAA
nCRPr.  CATAGGCACCCCAGGCTTGACACTTTATGCTTCCGGCTCGTATAATGTGTGCATGGATAA

pLAC.   GCGGATAACAATTTCACACAGGAAACAGCTATG
CRPr.   GTAGCTAGGAATTTCACACTGCAAACAGCTATG
nCRPr.  GTAGCTAGGAATTTCACACTGCAAACAGCTATG
```

*Figure S11:* Overview of promoter sequences used in this manuscript. The top row indicates the original LacZ upstream region including start codon ATG (NCBI; gene ID 945006, NC_000913.3), whilst the 2nd and 3rd row give the sequence of the engineered CRPr and nCRPr promoters. Colour indicates CRP binding sites according to [8] (yellow), [9] (green) or both (purple), and the LacI binding site (blue) according to [8]. In grey, changes in the engineered promoters are indicated.

### References

[1] D. J. Kiviet, P. Nghe, N. Walker, S. Boulineau, V. Sunderlikova, and S. J. Tans, "Stochasticity of metabolism and growth at the single-cell level," *Nature*, vol. 514, pp. 376–379, Oct. 2014.

[2] C. You, H. Okano, S. Hui, Z. Zhang, M. Kim, C. W. Gunderson, Y.-P. Wang, P. Lenz, D. Yan, and T. Hwa, "Coordination of bacterial proteome with metabolism by cyclic AMP signalling," *Nature*, vol. 500, pp. 301–306, Aug. 2013.

[3] J. W. Young, J. C. W. Locke, A. Altinok, N. Rosenfeld, T. Bacarian, P. S. Swain, E. Mjolsness, and M. B. Elowitz, "Measuring single-cell gene expression dynamics in bacteria using fluorescence time-lapse microscopy," *Nature Protocols*, vol. 7, pp. 80–88, Jan. 2012.

[4] N. Walker, *A single-cell study on stochasticity growth and gene expression.* Dissertation (TU Delft), Apr. 2016. ISBN: 9789492323026.

[5] I. T. Kleijn, L. H. J. Krah, and R. Hermsen, "Noise propagation in an integrated model of bacterial gene expression and growth," *PLOS Computational Biology*, vol. 14, p. e1006386, Oct. 2018.

[6] L. Susman, M. Kohram, H. Vashistha, J. T. Nechleba, H. Salman, and N. Brenner, "Individuality and slow dynamics in bacterial growth homeostasis," *Proceedings of the National Academy of Sciences*, vol. 115, pp. E5679–E5687, June 2018.

[7] B. D. Towbin, Y. Korem, A. Bren, S. Doron, R. Sorek, and U. Alon, "Optimality and suboptimality in a bacterial growth law," *Nature Communications*, vol. 8, pp. 1–8, Jan. 2017.

[8] J. M. Hudson and M. G. Fried, "Co-operative interactions between the catabolite gene activator protein and the lac repressor at the lactose promoter," *Journal of Molecular Biology*, vol. 214, pp. 381–396, July 1990.

[9] C. L. Lawson, D. Swigon, K. S. Murakami, S. A. Darst, H. M. Berman, and R. H. Ebright, "Catabolite activator protein: DNA binding and transcription activation," *Current Opinion in Structural Biology*, vol. 14, pp. 10–20, Feb. 2004.
